# Activating an adaptive immune response from a hydrogel scaffold imparts regenerative wound healing

**DOI:** 10.1101/2020.05.27.117317

**Authors:** Donald R. Griffin, Maani M. Archang, Chen H. Kuan, Westbrook M. Weaver, Jason S. Weinstein, An Chieh Feng, Amber Ruccia, Elias Sideris, Vasileios Ragkousis, Jaekyung Koh, Maksim V. Plikus, Dino Di Carlo, Tatiana Segura, Philip O. Scumpia

**Author notes:** Donald R. Griffin and Maani M. Archang contributed equally to this work. Tempo Therapeutics, San Diego, CA 92109. To whom correspondence should be addressed, Philip O. Scumpia, Tatiana Segura.

## Abstract

Biomaterial scaffolds represent a promising approach for material-based tissue regeneration. We previously developed microporous annealed particle (MAP) hydrogels - a flowable, microparticle-based hydrogel in which neighboring hydrogel particles are linked *in situ* to form a porous scaffold that accelerates wound healing. To promote more extensive tissue ingrowth before scaffold degradation, we aimed to slow scaffold degradation by switching the chirality of the crosslinking peptides from L-peptides to D-peptides. Unexpectedly, despite showing the predicted slower enzymatic degradation *in vitro*, D-peptide crosslinked MAP hydrogel (D-MAP) hastened material degradation *in vivo* and imparted significant tissue regeneration to healed cutaneous wounds, including increased tensile strength and hair neogenesis. By themselves, D-chiral peptides were poor activators of macrophage innate immune signaling *in vivo*, but MAP particles elicit IL-33 type 2 myeloid cell recruitment which is amplified *in vivo* in the presence of D-peptides. Remarkably, D-MAP elicited significant antigen-specific immunity against the D-chiral peptides, and an intact adaptive immune system was required for the hydrogel-induced skin regeneration. These findings demonstrate that the generation of an adaptive immune response from a biomaterial is sufficient to induce cutaneous regenerative healing despite faster scaffold degradation.

---

The ultimate goal of regenerative medicine is the restoration of tissue function back to physiologically normal activity. For hydrogel-based biomaterial scaffold strategies, this has led to a gold standard of balancing material degradation with endogenous tissue regrowth. The multitude of factors present in a clinical setting (e.g. age, relative health, etc.) leads to a wide variation in chemical and physical parameters *in situ*, which makes striking a robust degradative-regenerative balance particularly difficult. Our recent development of a flowable, granular biomaterial, known as MAP (Microporous Annealed Particle) gel, provides a new approach to make the balance more feasible (i.e. cellular ingrowth is decoupled from material degradation)^1^. The MAP gel is composed of microsphere building blocks, which imperfectly pack, leading to a continuous network of interconnected micrometer-scale void spaces that allows for the infiltration of surrounding tissue without the prerequisite degradation of material^1,2^. This unique interconnected void space design, termed open-porous architecture, resulted in improved tissue closure timelines, promotion of maturing vascular structures, and greatly diminished host immune response relative to a nanoporous (but chemical formulation equivalent) hydrogel in a cutaneous wound model^1^.

It is generally believed and reported that the ability of scaffolds to provide mechanical support to the growing tissue and promote wound healing is impacted by the degradation rate of the scaffold; too fast equates to loss of scaffold structure and too slow leads to fibrous encapsulation of the remaining material^3^. For MAP scaffolds, degradation leads to a slow loss of open-porous architecture and, subsequently, reduced tissue ingrowth. We hypothesized that modulating the degradation rate of MAP scaffolds would maintain the open-porous architecture and influence both wound closure rate and regenerated tissue quality. Our approach focused on prolonging MAP scaffold integrity, while maintaining hydrogel composition as close to its previous form as possible (chemically and physically).

Previously, we used amino acid chirality to tune the proteolysis rate of peptide nanocapsules to control release of encapsulated growth factors^4^. Building from this work, we chose to use an analogous approach of switching the chirality of the MAP peptide crosslinker targeted by enzymolysis *in vivo* (e.g. the site of matrix metalloprotease (MMP)-mediated bond cleavage). We hypothesized that this approach to modulating scaffold degradation rate would maintain the hydrogel micro-environment (hydrophobic-hydrophilic balance, mesh size, physical moduli, etc.), while also increasing the long-term hydrogel integrity to allow full infiltration of cells capable of laying new extracellular matrix, thus providing a greater integration of the entire construct with the host tissue.

As mentioned, changing the chirality of peptide moieties leads to a diminished degradation rate by endogenously present enzymes^4,5^, and has been used, specifically in hydrogel formulations, to control material properties. The use of chirality was made more attractive by the fact that polypeptides of D-enantiomeric amino acids are not reported to typically elicit a robust immune response and are considered poorly immunogenic or even possess tolerizing effects by themselves^5^. We expected this enantiomeric shift in crosslinker would allow us to build on the success of our previous studies by avoiding changes in material properties in exchange for some potentially modest immunomodulatory effects. Notwithstanding, while the innate and adaptive immune system undoubtedly are required to elicit the foreign body response and eventual fibrosis of some implanted biomaterial scaffolds^6,7^, recent evidence suggests that activation of the correct innate and adaptive immune responses is critical for enhancing the regenerative ability of a biomaterial scaffold^8,9^.

In the current study, we demonstrate that while hydrogels containing D-amino acid crosslinking peptides are more resistant to proteolytic degradation *in vitro*, they undergo more rapid clearance *in vivo* in a murine wound healing model. To our surprise, although incorporation of D-amino acid hydrogels resulted in faster degradation than L-amino acid hydrogels, they induced tissue regenerative response with hair follicle neogenesis and greater tissue tensile strength in fully healed wounds. We find that a subtle type 2 innate immune response triggered by the MAP hydrogel was amplified by adaptive immune activation against D-amino acids within the hydrogel and mice without an adaptive immune system did not have the amplification of the immune response and were unable to elicit skin regeneration with D-MAP. These findings provide additional evidence that direct activation of an appropriate adaptive immune response may be used to elicit regenerative healing.

## Incorporation of D-amino acids into the crosslinking peptides slows in vitro degradation of MAP hydrogel

For MAP scaffolding to maintain structural support throughout the wound closure process, we hypothesized that the scaffold degradation should occur after wound closure has been achieved. We thus used enantiomeric chemistry to change degradation rate without changing the initial material properties (e.g. hydrophobicity, mesh size, and charge) of the hydrogel^4^. All amino acids at the site of enzymatic cleavage for the MMP-degradable peptide were changed to D-amino acids (**see Methods** for peptide sequence). We matched the stiffness (i.e. storage modulus) by rheology of both the D-enantiomeric MAP (D-MAP) and L-enantiomeric (L-MAP) formulations to that used in our previous MAP-based cutaneous application (~500Pa; see Supplemental Figure 1a), which required minimal changes to peptide crosslinker content (L-MAP – 4mM; D-MAP – 4.1mM). After formulation optimization, we generated the microsphere particles using a previously published microfluidic technique^1^. Following application of Collagenase I to L-MAP, D-MAP or a 50% mixture of D-MAP and L-MAP (1:1 L/D-MAP), the L-MAP hydrogel degraded within minutes, while the degradation of D-MAP by itself or within a mixture with L-MAP was minimal even after an 1 hour, as expected (Supplemental Figure 1b and Figure 1a).

**Figure 1.**
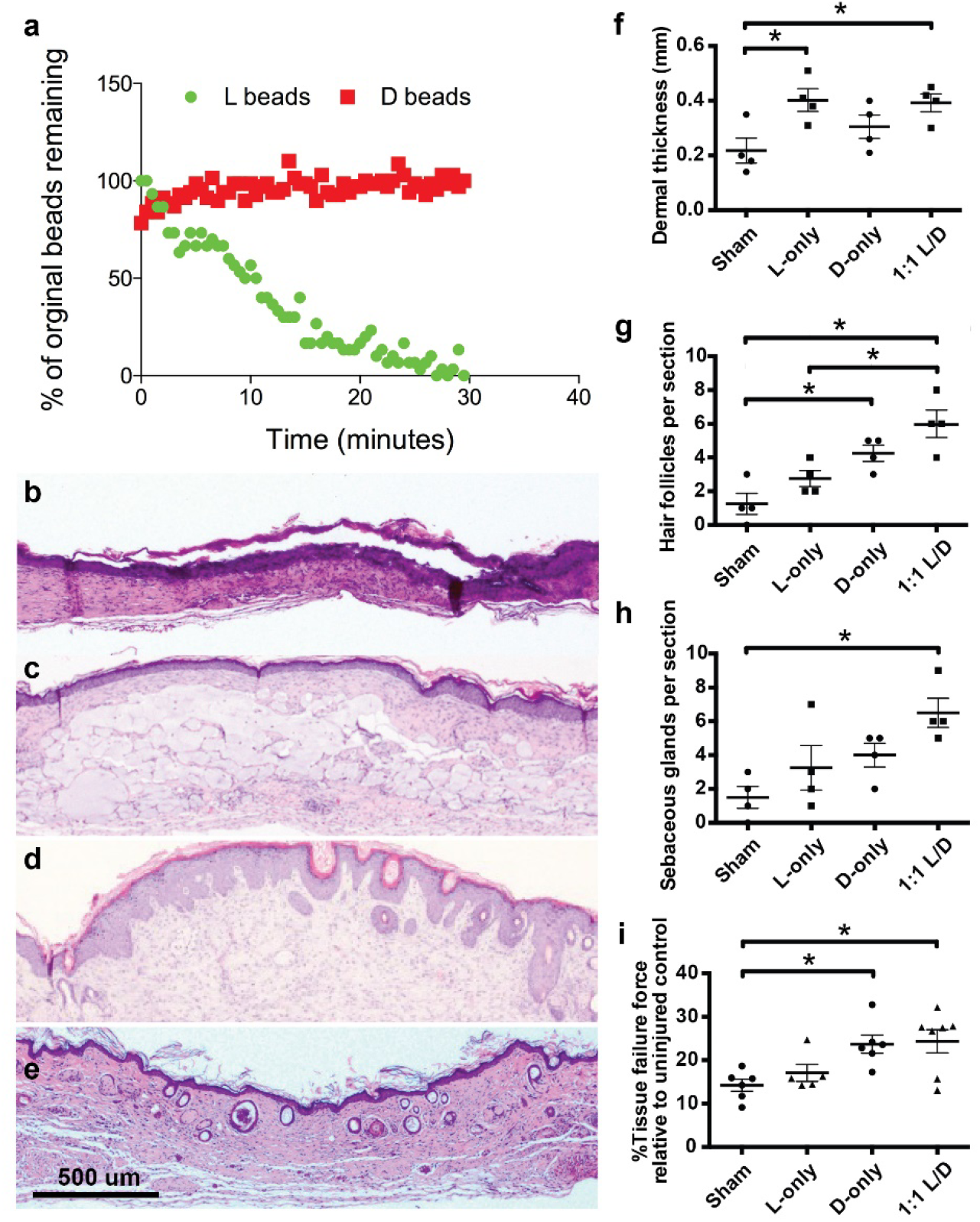
Presence of D-chiral crosslinker peptides decreases hydrogel degradation *in vitro* but enhances hydrogel degradation in SKH1 hairless mice. **a)** Fabricated L or D hydrogels were tested for *in vitro* enzymolysis behavior through exposure to a solution of collagenase I (5U/mL). **b-e)** Representative low power view of H&E sections from healed skin 21 days after splinted excisional wounding from a Sham (b), L-MAP (c), D-MAP (d), and 1:1 mixture of L-MAP and D-MAP treated wound in SKH1 mice (e). **f-h)** Histologic quantification of dermal thickness (in mm), hair follicles, sebaceous glands. Each point represents average of 2 sections from 2 separate slides of one wound. The dots in the plots represent one animal. **i)** 28 days after incisional, unsplinted wounds were created, healed wounds that were treated without or with different hydrogels were tested against unwounded skin in the same mouse. Tensile strength was evaluated by tensiometry and reported as a percentage of the tensile strength of the scar tissue when compared to the normal skin of the same mouse. Data is plotted as a scatter plot showing the mean and standard deviation. *, **, *** represent statistical significance by ANOVA using the Tukey multiple comparisons test to determine statistical significance.

## Incorporation of D-amino acids accelerates MAP degradation in murine wounds

We next examined how D-MAP compares to L-MAP *in vivo* in a murine splinted excisional wound model^1,10^. We did not find any difference in wound closure rate, or any increased erythema or gross signs of inflammation in wounds treated with D-MAP, L-MAP, or a 1:1 mixture of L/D MAP any time after treatment (shown Day 3 and Day 6 after wounding, Supplemental Figure 2a). When comparing wound closure to sham treatment (no hydrogel), we found that 50% mixture of L/D MAP induced more rapid wound closure (assessed on Day 9 after wounding) than sham (Supplemental Figure 2b), similar to previous results with L-MAP hydrogel^1^. Seeing no differences in wound closure results were noted when D-chiral cross-linking peptides were incorporated into the hydrogel in comparison to L-chiral cross-linked hydrogels, we next examined whether the degradation of hydrogels containing D amino acid cross-linkers was slowed *in vivo* by examining excised tissue 21 days after the wound was completely healed. As expected, sham treated mice displayed no hydrogel (Figure 1b). Unexpectedly, while histologic sections of wounds treated with L-MAP hydrogel displayed significant hydrogel persistence 21 days after wounding in various stages of degradation, those wounds treated with D-MAP or a 1:1 L/D-MAP displayed minimal to no hydrogel remaining at the wound site (Figure 1c-e).

**Figure 2:**
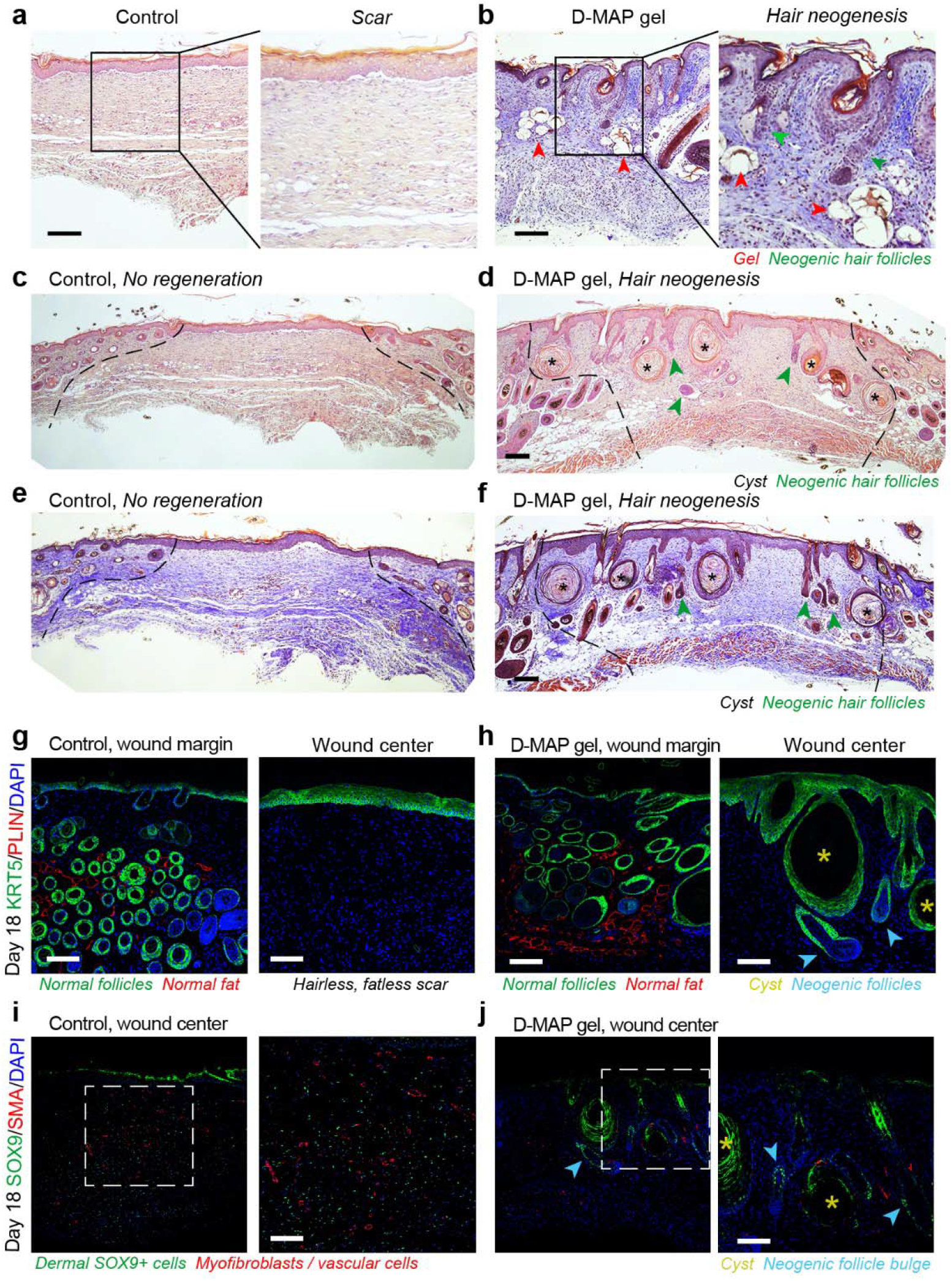
D-MAP hydrogel induces neogenesis of hair follicles in full-thickness skin wounds in B6 mice. **a-f)** H&E (a, c, d) and Trichrome staining (b, e, f) of healed 4-mm full-thickness splinted skin wound on day 18. Control (sham-treated) wounds heal with scarring (a, c, e), while D-MAP gel treated wounds form numerous epidermal cysts (asterisks) and, prominently, regenerate de novo hair follicles (green arrowheads) (b, d, f). In some instances, neogenic hair follicles form in close association with epidermal cysts. As compared to normal, pre-existing anagen hair follicles at the wound edges, neogenic hair follicles display early anagen stage morphology (Wound edges in b-d are outlined and D-MAP hydrogel remnants in b are marked with red arrowheads). **g-h)** Immunostaining for epithelial marker KRT5 (green) and adipocyte marker PLIN (red), reveals normal KRT5^+^ anagen hair follicles and many mature PLIN^+^ dermal adipocytes (left panels in g and h). Regeneration of new KRT5+ hair follicles (arrowheads in h) along with KRT5+ epidermal cysts is observed only in D-MAP hydrogel-treated wounds (right panel g vs. h). No neogenic adipocytes are observed in hair-forming D-MAP-treated wounds. **i-j)** Immunostaining for SOX9 (green) and SMA (red), reveals many SOX9^+^ epithelial cells within the bulge region of neogenic hair follicles in day 18 DMAP-treated wounds (arrowheads in j). In contrast, in control (sham-treated) wounds that undergo scarring, dermal wound portion contains many Sox9^+^ cells, many of which also co-express contractile marker SMA (i). Expression of SMA is also seen in both control and D-MAP-treated samples in blood vessels. Scale bars, **a-j** – 100 μm.

## Presence of D-chiral cross-linking peptides in MAP hydrogels imparts tissue regenerative properties

Of note, initial examination of histologic sections of D-MAP and 1:1 L/D-MAP displayed a much different overall appearance than that of healed sham- or L-MAP-treated wounds. Previous reports suggest that, unlike large excisional wounds in adult mice (wounds larger than 1×1 cm) that result in significant regenerative healing with wound induced hair neogenesis (WIHN)^11–13^, wounds smaller than 1×1cm in mice, like the 4-mm punch biopsies performed in our studies, typically heal without regeneration of new hair and fat and, instead form scars^12,14,15^. Despite these reports, through transgenic activation of specific Hedgehog signals, it has been shown that when the correct regenerative cues are provided from wound fibroblasts, small wounds are capable of undergo regeneration^16^. Consistent with these results, histological examination of 4-mm excisional splinted wounds in mice that did not receive hydrogel (sham) displayed the typical appearance of scar tissue with a thinned, flattened epidermis, a thinned dermis with horizontally-oriented collagen bundles, vertically-oriented blood vessels, and lack of hair follicles and sebaceous glands (Figure 1b, f-h). Tissue from mice treated with L-MAP hydrogel displayed a similar flattened epidermis, but a thickened dermis compared to sham wounds (Figure 1f). This finding is not surprising given the substantial residual hydrogel remaining. Within the dermis surrounding the hydrogel, we observed fibroblasts secreting collagen/extracellular matrix and blood vessels forming between the hydrogel microparticles (Figure 1c). Only rare hair follicles and associated sebaceous glands were observed in the previously wounded areas of L-MAP treated tissue (Figure 1c, g-h). Remarkably, however, examination of histological sections of the D-MAP or 1:1 L/D-MAP treated tissue revealed a *de novo* regenerated appearance. The overlying epidermis often displayed undulation, while the dermis was thicker with a less parallel appearance of collagen fibers, and numerous immature looking hair follicles were seen spanning the length of the healed full thickness injury (Figure 1d-h). Like samples treated with L-MAP hydrogel, those treated with D-MAP or 1:1 L/D-MAP displayed increased skin thickness despite less hydrogel remaining in these samples (Figure 1f). Many samples also displayed epidermal cyst formation. In samples that displayed residual hydrogel, hair follicles were apparent directly overlying the degrading MAP hydrogel particles (Supplemental Figure 2c). The presence of hair follicles in SKH1 mice was suggestive of embryonic-like tissue regeneration, a phenomenon not often in the murine small wound model.

To better quantify tissue regeneration, we next performed tensile strength testing on unsplinted incisional wounds in SKH1 mice using a modified literature protocol^17^ to determine whether improved tensile strength was observed in any of the MAP-treated wounds over the sham treated wounds. We found that scar tissue from sham wounds revealed tensile strength that was approximately 15% of unwounded skin from the same animal (Figure 1i). While treatment of wounds with L-MAP hydrogel did not result in a significant increase in tissue tensile strength, treatment with either D or L/D MAP resulted in an ~80% improvement in tensile strength (Figure 1i). Taken together with regenerating hair follicles and increased dermal thickness, these findings highlight the ability of D-MAP containing hydrogels to elicit tissue regeneration.

## The hair follicles in D-MAP-treated wounds are neogenic

We next repeated the wound healing experiments in B6 mice to investigate if the regenerative phenomenon observed in D-MAP treated wounds was consistent with it being wound induced hair neogenesis (WIHN). We chose sham as control and D-MAP as a treatment method that showed evidence of regeneration in SKH1 mice. Similar to the sham and L-MAP treated wounds in SKH1 mice, the B6 mice wounds without hydrogel (sham) displayed the typical scar tissue with flat epidermis, thinned dermis with horizontally-oriented collagen bundles, vertically-oriented blood vessels, and lack of hair follicles and associated sebaceous glands and were confirmed by Masson Trichrome staining (Figure 2a, c, e). In contrast, histological sections of the D-MAP treated tissue revealed clear signs of WIHN. As in SKH1 mice, D-MAP treated B6 mice wounds displayed undulations and numerous epidermal cysts under the epidermis, while the dermis was thicker with a less paralleled dermal collagen fibers. Importantly, many neogenic hair follicles developed in the wound (Figure 2b, d, f). The neogenic hair follicles were in early anagen phases with immature appearance, yet many of them already had formed new sebaceous glands (Figure 2b) and featured prominent SOX9^+^ bulge stem cell region (Figure 2j). In several instances, neogenic hair follicles were physically connected to epidermal cysts (morphology not expected from pre-existing follicles). This suggests that in D-MAP treated wounds, epidermal cysts can be the initiation sites for de novo morphogenesis for at least some of the neogenic hair follicles (Figure 2h). Masson trichrome staining confirmed the presence of neogenic hair follicles within the collagen matrix of the wound bed (Fig 2b, f). Furthermore, regenerating day 18 D-MAP treated wounds with neogenic hair follicles lacked PLIN^+^ dermal adipocytes (Figure 2h), which is consistent with the fact that neogenic adipocytes in large wound-induced WIHN regenerate only once the neogenic hair follicles have matured, on week four after wounding^18,19^. Thus, addition of D-MAP to normally non-regenerating 4-mm excisional wounds activates neogenesis of hair follicles.

## D-MAP hydrogel enhances myeloid cell recruitment following subcutaneous implantation

Since the inclusion of D-amino acids led to an unexpected enhanced/earlier degradation of MAP hydrogel *in vivo* despite slowing the degradation kinetics in response to direct treatment with MMPs *in vitro*, we hypothesized that the D-amino acids in the hydrogel resulted in a degradation-facilitating environment through protease activation during an immune response.

To determine whether D-MAP can elicit an immune response in tissue in the absence of an inflammatory wound healing response, and to ensure sufficient hydrogel is present 21 days after implantation, we utilized a subcutaneous implantation model which also allows for larger amounts of hydrogel to be implanted than in the excisional wound model. We observed that hydrogel was present in subcutaneous implants of L, D, and 1:1 L/D-MAP hydrogels 21 days after subcutaneous implantation. To test whether the subcutaneous implants of D-MAP hydrogel resulted in enhanced immune cell recruitment, we utilized immunofluorescent microscopy with AlexaFluor488 labeled MAP hydrogel. We found that implants containing just L-MAP display only a background level of CD11b cells within the hydrogel, as previously observed^1^, while D-MAP or L/D-MAP resulted in the accumulation of CD11b-expressing myeloid cells within and around the scaffold (Figure 3a-b). These studies suggest that the presence of D-amino acids within the hydrogel results in myeloid cell recruitment to the hydrogel.

**Figure 3.**
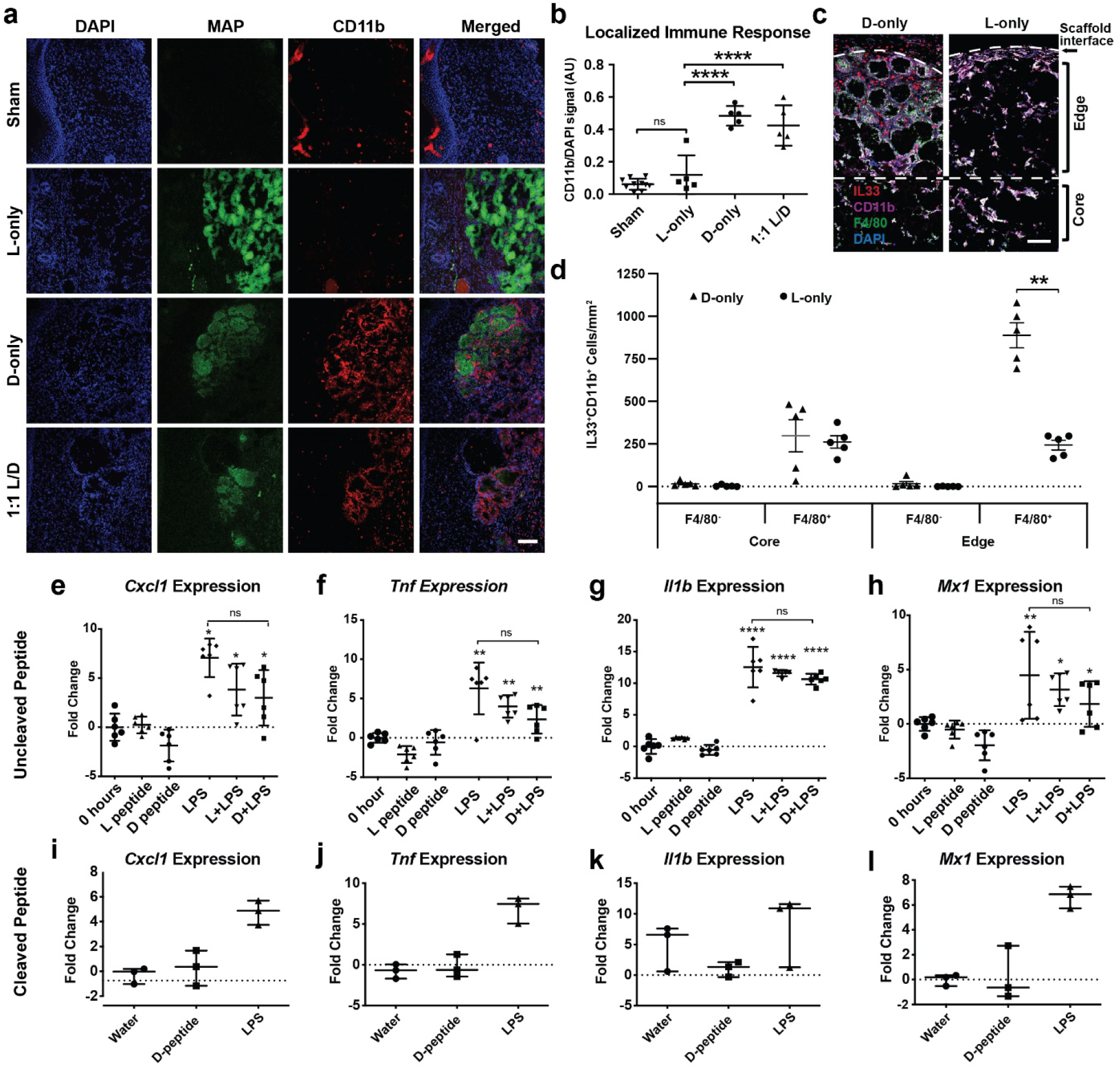
Myeloid cell activation and accumulation is associated with increased clearance of hydrogel containing D-chiral peptides, but D-chiral peptides do not directly induce transcriptional activation of myeloid cells *in vitro* through pattern recognition receptors engagement. **a)** Representative confocal immunofluorescent images staining myeloid cells (CD11b^+^) within healed wounds of B6 mice in the absence or presence of the indicated hydrogel. Scale = 100μm. **b)** Quantification of CD11b^+^ cellular infiltrate in healed tissue 21 days after wounding in the presence or absence of hydrogel. **c-d)** Representative high-resolution confocal immunofluorescence imaging for CD11b, F4/80, DAPI, and IL-33 from subcutaneous implants of L- or D-MAP hydrogel implants (c) and quantification of IL-33 producing macrophages and other myeloid cells at hydrogel edge and core (n=5) (d). **e-h)** Murine bone marrow derived macrophages (BMDMs) from B6 mice were stimulated with 500 μg/ml of full-length L- or D-crosslinker peptide in the presence or absence of lipopolysaccharide (10ng/ml) for 6 hours. Shown are qPCR results of 4 inflammatory genes from two separate experiment performed with n = 3. **i-l)** BMDMs were stimulated with LPS (10ng/ml) or cleaved D-crosslinker peptide (500μg/ml) that possessed an N-terminal D-amino acid. Experiment was performed in triplicate. Data is plotted as a scatter plot showing the mean and standard deviation. *, **, *** represent statistical significance by ANOVA using the Tukey multiple comparisons test to determine statistical significance.

To further characterize how D-MAP enhances innate immune recruitment, we next evaluated histology of subcutaneous implants. After 21 days, L- or D-MAP implants were removed from the mice. As we previously demonstrated^20^, most L-MAP implants did not display traditional fibrous capsule formation and only a few areas displayed some focal and thin fibrous capsules containing small foci of immune cells (Supplemental Figure 3a). No foreign body reaction with giant cells were noted within or surrounding the hydrogel, but some lymphohistiocytic immune infiltrates were seen dispersed randomly within and in areas towards the edge of the implant (Supplemental Figure 3a). This is likely a property of the hyperporous nature of the MAP hydrogel allowing cells to migrate through the hydrogel rather than typical materials that allow a build-up of immune cells at the edges of the hydrogel. In contrast, D-MAP implants displayed thick fibrous capsules filled with mainly lymphohistiocytic cells and rare scattered neutrophils and eosinophils (Supplemental Figure 3a). Epithelioid histiocytes degrading hydrogel particles with scattered surrounding lymphocytes, and rare eosinophils and neutrophils were detected at the edges of the scaffold. Many more of these immune aggregates were found within D-MAP scaffolds than L-MAP scaffolds (denoted by *s in Supplemental Figure 3a). These findings are not consistent with a typical type 1 foreign body granuloma (no foreign body giant cells with sheaths of surrounding lymphocytes) associated with a strong Th1/M1 immune response seen in tuberculosis or with inflammatory materials. Instead, the material was eliciting a type II foreign body reaction typically associated inert foreign materials such as silicone^21–23^. Immunofluorescent staining for F4/80 and CD11b confirmed the increased presence of F4/80^+^CD11b^+^ macrophages within the D-MAP scaffolds at 21 days with concentrations at the edges of the hydrogel (Supplemental Figures 3b and 3c). These data suggest that D-MAP either possesses increased adjuvant properties by itself, or the MAP hydrogel itself that is being amplified by the presence of D-chiral peptides or the MAP hydrogel possesses adjuvant properties and the presence of D-chiral peptides induce an adaptive immune response that can dramatically amplify the immune response to the MAP hydrogel.

Allergic responses and parasites can elicit a type 2 immune response including atypical type 2 granulomatous responses at least partially through IL-33 production by epithelial cells, recruited myeloid cells and resident macrophages^21,22,24,25^. A recent study uncovered that implanted, non-degradable microparticle-based materials elicit an interleukin (IL)-33-dependent type 2 innate immune response form circulating CD11b^+^ myeloid cells and macrophages^26^. It is possible that MAP particles could activate this same program, especially given the atypical type 2 foreign body responses observed in D-MAP samples. Indeed, 21 days after implantation, we found similar numbers of IL-33-expressing F4/80^+^CD11b^+^ macrophages in the center/core of both L- and D-MAP implants (Figure 3c and 3d), consistent with both L- and D-MAP samples activating this type 2 pathway. In addition, there was a dramatic increase in IL-33 producing IL-33^+^F4/80^+^ macrophages at the edges of D-MAP implants, when compared to L-MAP implants, suggesting that amplification of this response was indeed occurring when a potential antigen are present (Figure 3c and 3d). These results confirm that the hydrogel possesses a type 2 innate “adjuvant” effect, that can activate the adaptive immune system, and may contribute to immune activation with D-MAP hydrogel. When native L-peptides are cross-linking the MAP scaffolds, the immune response remains mild as the hydrogel degrades slowly over time^20^, but the presence of D-peptide accelerates immune mediated degradation by possibly activating an adaptive immune response. Therefore, the D-peptides may possess additional adjuvant effects over the MAP hydrogel alone or may be acting as an antigen in the presence of the mild adjuvant effect of MAP to augment immune responses to D-MAP.

## D-amino acid containing peptides do not directly induce traditional pathogen recognitional receptor (PRR) inflammatory gene expression in macrophages

We next tested whether D-peptides could directly activate innate immunity through traditional PRR-induced transcriptional response. We stimulated murine bone marrow derived macrophages (BMDMs) with full-length D-amino acid or L-amino acid containing peptides in the presence or absence of bacterial lipopolysaccharide, the Toll-like receptor 4 agonist that results in rapid macrophage transcriptional responses. We chose to examine genes that are reliably and strongly inducible downstream of the major signaling pathways to simultaneously interrogate multiple PRR pathways and general macrophage activation pathways downstream of a variety of cellular insults (AP-1, MAPK, NF-κB, and type I IFN)^27–29^. To our surprise, neither L- or D-amino acid containing crosslinking peptides alone at high doses (1mg/ml) induced the expression of pro-inflammatory genes *Tnf* (NF-κB dependent), *Il1b* (NF-κB and MAPK dependent), *Cxcl2* (AP-1 dependent early response), or *Mx1* (type I IFN dependent) in murine BMDMs at 6 hours (t_max_ of gene induction; Figure 3e-l). Additionally, neither L- nor D-peptides enhanced the ability of LPS to induce the expression of these same genes (Figure 3e-h).

Previous studies have shown that peptides containing an N-terminal D-methionine can activate the innate immune receptor formyl peptide receptor 2 and formyl peptide like receptor 2^30–32^. Since cleavage of D-amino acid peptide can result in shorter peptides that contain a D-amino acid at the N-terminus, we next wished to examine whether a peptide corresponding to the cleaved D-peptide could activate inflammatory responses in BMDMs. Similar to the results with intact D-peptide, high concentrations of cleaved D-peptide (1mg/ml) did not induce the transcription of *Tnf*, *Il1b*, *Cxcl2*, or *Mx1* at 6 hours (Figure 3i-l). Since there is a very low likelihood that cleaved D-peptide will be present at such high local concentrations within the implanted hydrogel while it is being degraded *in vivo*, these findings argue that D-chiral peptides are poor activators of a traditional PRR-mediated inflammatory response in macrophages and suggest that D-peptides may be acting as antigens rather than adjuvants to enhance immunity, leading to enhanced degradation of D-MAP.

## D-MAP elicits antigen-specific humoral immunity

We next evaluated whether the D-MAP activates adaptive immunity. The adaptive immune system recognizes non-self-peptide antigens to induce cell mediated (T cell) and humoral (B cell) immunity. Peptides containing D-amino acids have been reported to activate or suppress T cell dependent and T cell independent adaptive immune responses^5,33^. In the context of the MAP, cross-linking peptides that are non-native may be presented to the immune system until fully degraded. . D-peptides could be presented by antigen presenting cells directly to T cells, eliciting a T cell dependent adaptive immune response or alternatively, the presence of D-amino acid containing peptides on the surface of a large molecule of MAP hydrogel could directly crosslink the B cell receptor, leading to T cell independent antibody responses similarly to the T cell independent antigens, like nitrophenyl-ficoll. To test the hypothesis that D-amino acids modulate adaptive immunity, we examined whether mice that were wounded or received subcutaneous implants of L-MAP, D-MAP, or 1:1 L/D-MAP were able to develop T-helper cell dependent (IgG1 or IgG2a) or T cell independent (IgG3) antibodies against L- or D-amino acid containing crosslinkers^34–37^.

Indeed, regardless of whether D-containing MAP hydrogel was applied to wounded tissue or given via subcutaneous implants, mice developed a T cell dependent IgG1 and IgG2a response against the D-amino acid containing peptide, but not a T cell independent IgG3 response. These results are more consistent with a T cell-dependent immune response against D-peptides (Figure 4a-b). IgG1 is typically associated with a Th2 “tissue repair” type response, while IgG2a is typically associated with a Th1 “foreign body” response that typically requires strong adjuvants to develop, depending on the strain of mice^38,39^. The fact that anti-D peptide-specific IgG2a induced when the hydrogel was given to mice in a wound environment but not when the hydrogel was given in the subcutaneous implant model suggests that, by itself, the hydrogel does not possess sufficient adjuvant effects to induce robust Th1 responses. However, the inflammation present in the wound environment may result in a mixed Th2/Th1 response to the D-MAP (Figure 4b and 3e). Mice that were treated with L-MAP alone did not develop antibody responses to L-peptide, but interestingly, mice that were treated with L- and D-MAP hydrogel developed antibodies against both L- and D-peptides (Figure 4e, 3h), suggesting that cross-priming and adaptive immune activation against L-peptide may occur when D-peptides are present.

**Figure 4.**
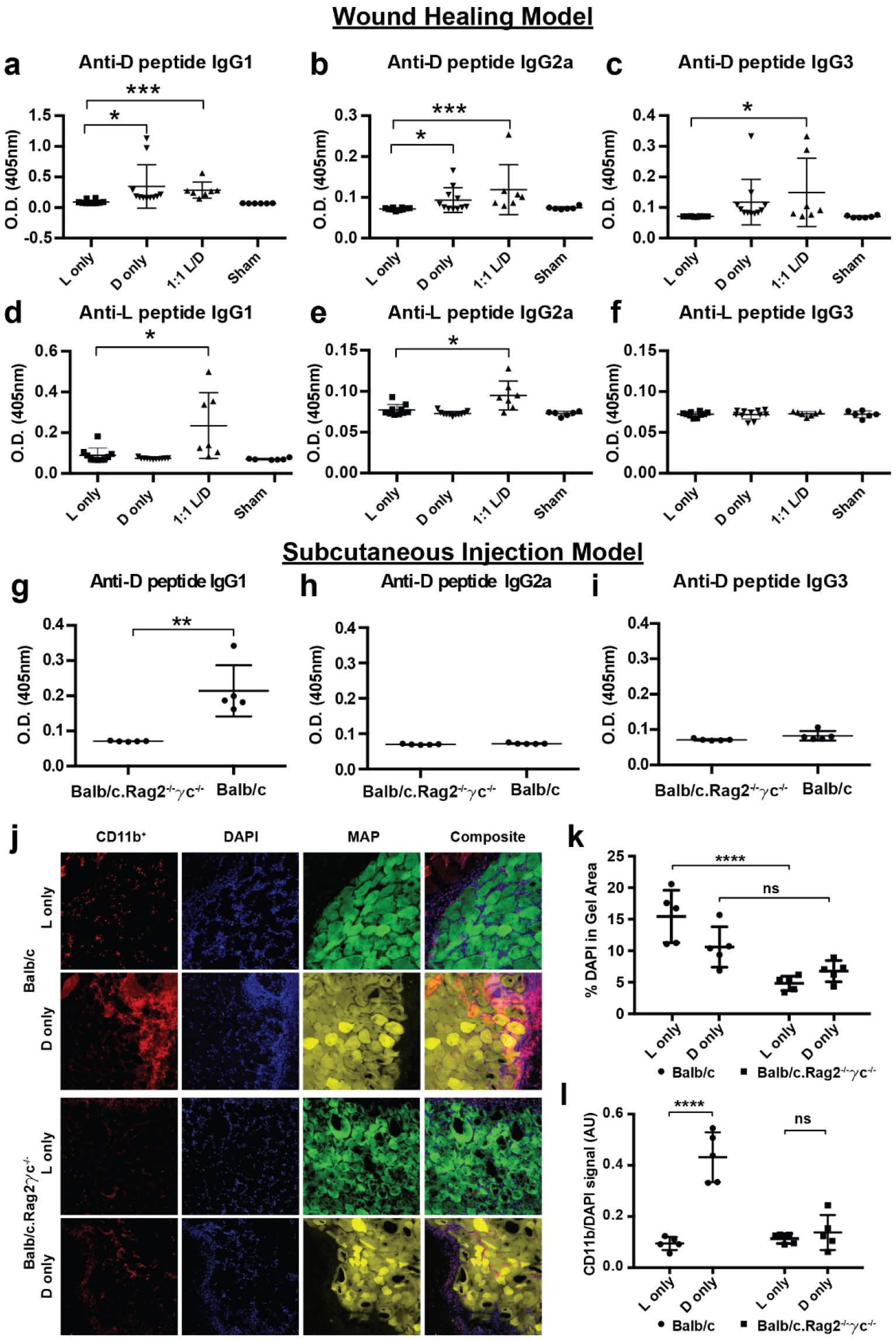
D-MAP induces antibody responses *in vivo* and requires an intact adaptive immunity for optimal myeloid cell recruitment. **a-c)** Measurement of anti-D specific IgG subtype antibodies by ELISA 21 days following wound healing experiments in SKH1 mice treated with indicated hydrogels. **d-f)** Measurement of anti-L specific IgG subtype antibodies by ELISA 21 days following wound healing experiments in SKH1 mice treated with indicated hydrogels. **g-i)** Measurement of anti-D specific IgG subtype antibodies in Balb/c or Balb/c.Rag2^−/−^γc^−/−^ mice given a subcutaneous injection of D-MAP 21 days after injection. **j-l)** Representative examples of confocal immunofluorescent imaging for CD11b, DAPI, and hydrogel from subcutaneous implants of L- or D-MAP hydrogel implants in Balb/c or Balb/c.Rag2^−/−^γc^−/−^ mice (j) and quantification of total DAPI+ cells (k), CD11b^+^ myeloid cells (l). Data is plotted as a scatter plot showing the mean and standard deviation. *, **, *** represent statistical significance by student t-test for the comparison indicated.

## CD11b myeloid cell recruitment to D-MAP requires an intact adaptive immune system

Our data suggest that the activation of adaptive immune responses to D-MAP contributes to immune infiltration and degradation of D-MAP. To test this hypothesis further, we examined whether Balb/c.Rag2^−/−^γc^−/−^ mice, which are devoid of an adaptive immune system and innate lymphoid cells, but possess a fully functional myeloid system^40^. Indeed, the recruitment of CD11b^+^ myeloid cells to D-MAP hydrogel in Balb/c.Rag2^−/−^γc^−/−^ mice was decreased to comparable levels to those seen in L-MAP in WT mice (Figure 4l). Total cellularity within the hydrogel in Balb/c.Rag2^−/−^γc^−/−^ mice was also diminished to D-MAP hydrogels in Balb/c.Rag2^−/−^γc^−/−^ mice (Figure 4l). The adaptive immune system is required for D-MAP-induced skin regeneration.

Since we found that MAP hydrogel annealed with D-amino acids requires adaptive immunity to induce immune cell recruitment, we next wished to determine whether D-MAP requires an adaptive immune system to induce hair follicle regeneration. Further, while we found D-MAP increases tensile strength in B6 mice, our initial findings of WIHN were obtained in hairless SKH1 mice, a mouse that can only develop vellus hairs. We next tested whether D-MAP can induce WIHN in B6 mice, a mouse that develops terminal hairs. To this end, we performed excisional splinted wounds in B6 and B6.Rag1^−/−^ mice untreated (sham) or treated with 1:1 L/D-MAP gel and examined them 25 days after wounding. Of note, because in preliminary studies, scars induced by 4-mm punch wounds healed with extremely small scars in B6 mice, we used a 6-mm punch in this experiment.

Sham wounds in B6 mice demonstrated an obvious depigmented, irregularly-shaped scar, while scars in B6 mice treated with 1:1 L/D-MAP gel were difficult to identify visually as they displayed hair growth over the wounds and less atrophy/surface changes typically seen in scars (representative example Figure 5a, all wound images Supplemental Figure 4). Scars in sham-treated or 1:1 L/D-MAP-treated B6.Rag1^−/−^ mice were smaller than those in sham-treated B6 mice, but were identifiable in B6.Rag1^−/−^ mice regardless of whether wounds were sham treated or hydrogel treated (Figure 5a). All wound (including 1:1 L/D-MAP-treated B6 wound areas) injuries were confirmed by examining the defect on the fascial side of the tissue after excision of skin for histological processing. Histological sections of the healed skin of mice displayed significant WIHN, including hair follicles and sebaceous gland formation only in wounds of wildtype mice treated with 1:1 L/D-MAP (Figure 5b-d, and Supplemental Figure 5). However, both sham wounds in B6 and Rag^−/−^ mice, and in the 1:1 L/D-MAP treated B6.Rag1^−/−^ mice displayed the presence of scars, without significant WIHN, confirming the requirement of the adaptive immune system in skin regeneration induced by D-peptide containing MAP gel (Figure 5b-d). These studies highlight that terminal hairs and complete adnexal structures (hair follicles with sebaceous glands) can be regenerated through adaptive immune activation by MAP hydrogel scaffolds, and further highlight a critical role of the adaptive immune system in the modulation of tissue regeneration.

**Figure 5.**
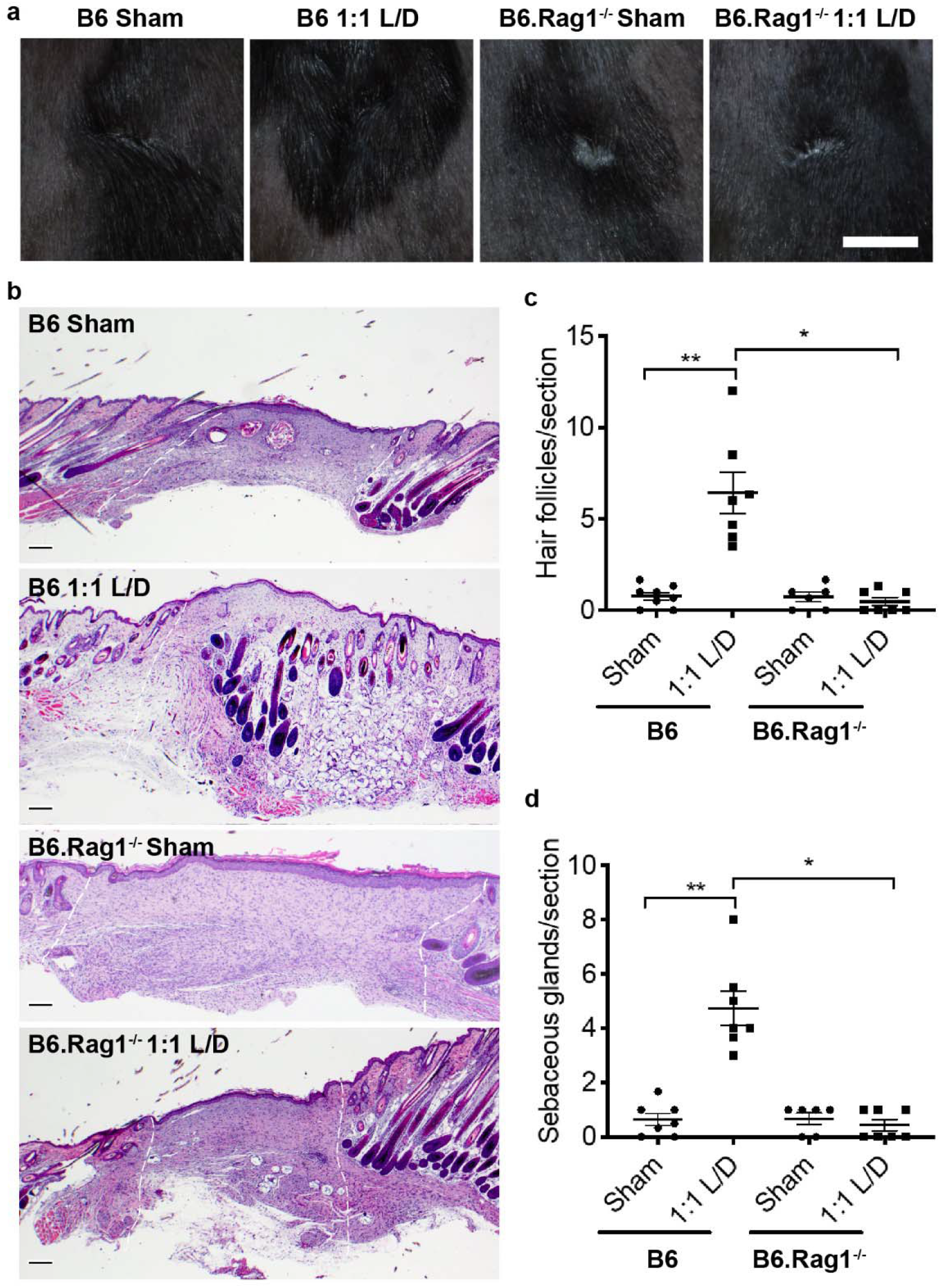
D-MAP requires an adaptive immune system to induce hair follicle neogenesis. **a)** Representative examples of images of healed splinted excisional wounds in B6 or B6.Rag1^−/−^ mice 17 days later treated as sham (no hydrogel) or 1:1 L/D-MAP treatment. Scale = 5mm. **b)** Histologic sections of healed tissue from B6 or B6.Rag1^−/−^ mice. Scale = 200μm. White dashed lines denotes wounded area. Enumeration of the average numbers of **c)** hair follicles (* denotes p=0.002, ** denotes p<0.001 by unpaired t test). **d)** sebaceous glands from 3 histological sections per sample from B6 mice and B6.Rag1^−/−^ mice. Data is plotted as a scatter plot showing the mean and SEM. * denotes p=0.0006 by Mann Whitney U test, ** denotes p<0.0001 by unpaired t test.

## Discussion

In most mammals, the natural process of scar formation and tissue fibrosis is highly evolved and is a tissue-scale attempt to restore critical barrier functions for survival. This process, however, is ultimately a biological ‘triage’ that favors the rapid deposition of a fibrotic matrix to restore the barrier at the expense of a loss of function of complex tissue. In the skin, this fibrotic response results not only in a loss of functioning adnexal structures, but skin tissue that is more fragile and prone to re-injury. A major goal of engineering skin regeneration is to allow for the rapid restoration of barrier function while providing increased tissue tensile strength and higher tissue function. Many biomaterial-based approaches have been attempted including addition of growth factors, decellularized extracellular matrix constructs, and stem cell therapeutics, and have seen limited success in the ability to restore function in wounds of various tissue. We previously showed that our synthetic MAP scaffold can accelerate wound closure in murine wounds^1^. Our findings resported here further highlight that incorporation of a modest adaptation to MAP, using D-enantiomer amino acids within the cross-linking peptides of MAP microparticle gels, induced two hallmarks of skin regeneration: hair neogenesis and improved tensile strength. This response was shown to be dependent on the generation of an adaptive immune response to D-amino acid containing peptides, and occurred without addition of stem cells, growth factors, or adjuvants.

While the activation of adaptive immunity can contribute to fibrosis, traditional foreign body granuloma formation, and rejection (i.e. loss of function) of biomaterial scaffolds or devices implanted into the body^6–8^, recent studies with growth factor containing extracellular matrices suggest that adaptive immune activation can also contribute to muscle regeneration ^8,9^. Further recent work has highlighted the potential of other biomaterials created to directly activate specific components of the immune system to treat cancer as immunotherapy platforms^41,42^. In concert, these studies suggest that the role of the adaptive immune system in wound healing is significantly more complex than previously realized. Our findings suggest that induction of a type 2 immune response to sterile microparticle-based materials can be used to trigger regeneration to degradable particles rather than fibrosis which occurs when particles are non-degradable. Our findings further support a role of adaptive immune cells to restore tissue function, as even in the absence of added growth factors or cells, the generation of an adaptive immune response from MAP scaffold induced tissue regeneration with stronger tissue that contained adnexal structures. Finally, we display the potential of the MAP scaffold as a potent immunomodulatory platform. Identifying factors produced by the immune system that tip the balance towards promoting regeneration instead of eliciting a foreign body response may lead to novel therapeutic targets to promote tissue regeneration by biomaterials.

## Materials and Methods

### L-MMP and D-MMP MAP Hydrogel Formation

Microfluidic water-in-oil droplet generators were fabricated using soft lithography, as previously described^1^. To enable microgel formation, two aqueous solutions were prepared. One solution contained a 10% w/v 4-arm PEG-vinyl sulfone (20 kDa, JenKem USA) in 300 mM triethanolamine (Sigma), pH 8.25, pre-functionalized with 500μM K-peptide (Ac-FKGGERCG-NH2) (GenScript), 500 μM Q-peptide (AcNQEQVSPLGGERCG-NH2), and 1 mM RGD (Ac-RGDSPGERCG-NH2) (GenScript). The other solution contained an 8mM di-cysteine modified Matrix Metalloprotease (MMP) (Ac-GCRDGPQGIWGQDRCG-NH2) (GenScript) substrate with either all L-chirality amino acid residues for L-MMP microgels or D-chirality amino acid substitution of amino acids at the site of MMP-mediated recognition and cleavage (Ac-GCRDGPQ_D_GI_D_W_D_GQDRCG-NH2) for D-MMP microgels. The oil phase was a heavy mineral oil (Fisher) containing 0.25% v/v Span-80 (Sigma). Downstream of the pinching region, a second oil inlet with a high concentration of Span-80 (5% v/v) was mixed with the flowing droplet emulsion. Both aqueous solution flow rates used were 0.75 μL/min, while both oil solutions were flowed at 4μL/min. The mixture was allowed to react overnight at room temperature and purified by repeated washes with an aqueous buffer of HEPES buffered saline pH 7.4 and pelleting in a tabletop centrifuge at 18000 × g for 5 mins. Raw materials are purchased endotoxin free and the final hydrogels are tested for endotoxin levels prior to implantation.

### Generation of MAP scaffolds from building block μgels

Fully swollen and equilibrated building block μgels were pelleted at 18000 × g for five minutes, and the excess buffer (HEPES pH 7.4 + 10 mM CaCl2) was removed by aspiration. Subsequently, building blocks were split into aliquots, each containing 50 μl of concentrated building blocks. An equal volume of HEPES pH 7.4 + 10 mM CaCl2 was added to the concentrated building block solutions. Half of these are spiked with Thrombin (Sigma) to a final concentration of 2 U/ml and the other half spiked with FXIII (CSL Behring) to a final concentration of 10 U/ml. These solutions were then well mixed and spun down at 18000 × g, followed by removal of excess liquid with a cleanroom wipe (American Cleanstat).

Annealing was initiated by mixing equal volumes of the building block solutions containing Thrombin and FXIII using a positive displacement pipet (Gilson). These solutions were well mixed by pipetting up and down, repeatedly, in conjunction with stirring using the pipet tip. The mixed solution was then pipetted into the desired location (mold, well plate, mouse wound, etc.) or loaded into a syringe for subcutaneous injection. The microgel fabrication was performed under sterile conditions. Following particle fabrication, 20 ul of dry particles were digested in 200 ul digestion solution (Collagenase IV 200 U/ml+ DNase I 125U/ml) and incubated in 37 C for 30 min before testing. Endotoxin concentrations were determined with Pierce LAL Chromogenic Endotoxin Quantitation Kit (Thermo Fisher Scientific) following manufacturer’s instructions. Particle Endotoxin levels were consistently below 0.2 Endotoxin U/mL.

### Degradation with collagenase

Microgel degradability was confirmed with collagenase I. A 1:1 v/v mixture of microgels formed with MMP-D- or MMP-L-sensitive cross-linker was diluted in collagenase I to a final concentration of 5 units collagenase/mL. This mixture was added to a 1 mm PDMS well and briefly allowed to settle. Images of the microgels were taken near the bottom of the well every 30 seconds for 2 hours with a confocal microscope. Image analysis was carried out through a custom MATLAB script (script provided by Dr. Sasha Cai Lesher-Perez) and ImageJ. MATLAB was used to determine the number of intact microgel spheres in each image. The previously mentioned script was applied with a minimum droplet radius of 30 pixels, a maximum droplet radius of 50 pixels, and a sensitivity factor of 0.98 for channel-separated images. Then, ImageJ was used to determine the area fraction fluorescing for each channel and each image. The thresholding for each image was set to a minimum of 50 and a maximum of 255 and the fluorescing area fraction was recorded.

### Mouse excisional wound healing model (Protocol# 10-011, UCLA IUCAC)

Mouse excisional wound healing experiments were performed as previously described^1,10^. Briefly 10-week old female CRL-SKH^Hr/hr^ mice (Charles River Laboratories; n=6), or 10-week old female C57Bl/6 (B6) or B6.Rag1^−/−^ mice (Jackson Laboratories; n=4 twice) were anesthetized using continuous application of aerosolized isoflurane (1.5 vol%) throughout the duration of the procedure and disinfected with serial washes of povidone-iodine and 70% ethanol. The nails were trimmed and buprenorphine 0.05mg/ml) was injected intramuscularly. The mice were placed on their side and dorsal skin was pinched along the midline. A sterile 4 mm biopsy punch was then used to create 2 through-and-through wounds, resulting in four clean-cut, symmetrical, full-thickness excisional wounds on either side of the dorsal midline. A small amount of adhesive (VetBond, 3M, Inc.) was then applied to one side of a rubber splint (O.D. ~12mm; I.D. ~8mm) and the splint was placed centered around the wound (adhesive side down). The splint was secured with eight interrupted sutures of 5-0 non-absorbable Prolene. A second splint wrapped in Tegaderm (3M, Inc.) was attached to the initial splint via a single suture to act as a hinged cover to allow wound imaging while acting as a physical barrier above the wound bed. After addition of thrombin (2 U/ml) and 10mM CaCl_2_, the experimental material (20 μL of L-only MAP, D-only MAP, 1:1 v/v mixture of L-MAP and D-MAP in HEPES-buffered saline containing Factor XIII (10U/ml) and 10mM CaCl2, or no hydrogel) was then added to one of the wound beds randomly to ensure each hydrogel treatment was applied to the different regions of wounded back skin to limit potential for site-specific effects. Following treatment, a Tegaderm-coated splint was applied, and wound sites were covered using a self-adhering elastic bandage (VetWrap, 3M, Inc.). Animals were housed individually to prevent wound manipulation. At the culmination of the wound healing experiment (Day 21 or Day 25) the mice were sacrificed by isofluorane overdose and cervical dislocation and imaged with a digital camera. The skin was excised and processed via either paraffin embedding for H&E or OCT blocks for immunofluorescence.

### Evaluation of wound closure

Wounds were imaged daily to follow closure of the wounds. Each wound site was imaged using high-resolution camera (Nikon Coolpix). Closure fraction was determined as described previously^1^. Briefly, closure was determined by comparing the pixel area of the wound to the pixel area within the 10 mm center hole of the red rubber splint. Closure fractions were normalized to Day 0 for each mouse/scaffold sample. Investigators were blinded to treatment group identity during analysis.

### Wound Imaging

On the specified day after wounds were created, close up images of wounds were taken using a Cannon Powershot A2600 or a Nikon D3400 DSLR Camera with an 18-55mm Lens, and were cropped to the wound area but not manipulated further. For wound closure, area was obtained using ImageJ by a subject blinded to the treatment.

### Tissue collection

After wounds healed, mice were sacrificed on the indicated day after wounding, and tissue collected with a ~5mm margin around healed wound. The samples were immediately submerged in Tissue-Tek Optimal Cutting Temperature (OCT) fluid and frozen into a solid block with liquid nitrogen. The blocks were then cryo-sectioned by cryostat microtome (Leica) and kept frozen until use. The sections were then fixed with 4% paraformaldehyde in 1X PBS for 30 minutes at room temperature, washed with 1X PBS, and kept at 4°C until stained. For antibody production analysis, blood harvested via cardiac puncture to obtain serum for ELISA.

### Macrophage cell culture

Mouse bone marrow derived macrophages were generated as previously described previously^28^. Briefly, following euthanasia, hindlimbs were removed aseptically and bone marrow was flushed. Bone marrow cells were cultured in CMG-conditioned complete DMEM media for 6 days. Cells were then treated with intact L- or D-peptide in ultra-pure H_2_0 at the indicated concentration in the presence or absence of LPS (10ng/ml). Cleaved D-peptide (with an N-terminal D-amino acid) (W_D_GQDRCG-NH2) was also used when indicated. Cells were harvested at 6 hours following treatment and expression of cytokines and chemokines was examined by qPCR using specific primers as described previously^43^.

### Incisional wound model

10-week old female C57Bl/6 mice (Jackson Laboratories) were anesthetized with isofluorane as above. The dorsal and side skin was dehaired using electric clippers followed by Nair (Church and Dwight, Inc.), then disinfected with serial washes of povidone-iodine and 70% ethanol. The nails were trimmed to lower the incidence of splint removal, and buprenorphine was injected IM as above. An incisional 2cm × 1cm wound was made with a scalpel. Mice (5 per group) were randomly assigned to receive 50μL of L-MAP, D-MAP, 1:1 v/v mixture of L-MAP and D-MAP or no hydrogel (aquafor). The mice were wrapped with Tegaderm followed by VetWrap as above.

### Histology and analysis

Samples were sectioned (6-10μm thick), then stained with hematoxylin and eosin or Masson trichrome by the UCLA Tissue Procurement Core Laboratory using standard procedures. Sections were examined by a board certified dermatopathologist (P.O.S.) and/or an expert in hair follicle neogenesis/regeneration (M.V.P.) who were blinded to the identity of the samples, for the presence of adnexal structures in tissue sections and dermal thickness. For enumeration, two to three tissue sections from the tissue block of each wound were examined and averaged per wound to obtain the count for each sample.

### Tensiometry

To evaluate the tensile properties of the healed incisional wounds, tensile testing was performed on an Instron model 3342 fitted with a 50N load cell and data recorded using the Instron Bluehill 3 software package. Tissue was collected from the wound site 28 days following wounding/treatment as a 2 cm × 4 cm “dumbbell” shape (with 1 cm center width in the handle portion). The sample was oriented such that the healed wound spanned the entire middle section of the dog bone (the thinner 1cm region) and the healed wound long axis was orthogonal to the direction of tension applied. The tissue sample was loaded into the Instron and secured with pneumatic gripers, pressurized to 40 PSI. The tissue was subjected to tensile testing at an elongation rate of 5 mm/min and ran through material failure.

For each tissue sample, stress/strain curves were calculated from force/elongation curves (provided from the Instron Bluehill software) using the known cross-sectional dimensions of the “dog bone” samples (each measured with calipers prior to placement on the Instron), and by measuring the starting distance between pneumatic grips with a caliper. The starting distance was standardized by preloading the sample to 0.5N, followed by measurement and then running of the tensile test to failure. This analysis enabled calculation of Yield Stress, which are reporting in Figure 1i.

### Subcutaneous implants of hydrogel

For subcutaneous implants, after anesthesia, 10-week old female Balb/c and Balb/c.Rag2^−/−^γc^−/−^ mice were injected with 50μL of L-MAP, D-MAP, or 1:1 v/v mixture of L-MAP and D-MAP (n = 5). 21 days later the skin and subcutaneous tissue containing the hydrogels were removed and processed for histology and immunofluorescence, and blood was collected by cardiac puncture to obtain serum for ELISA. B6 mice were used in another batch of experiments for immunofluorescence analysis and histology of subcutaneous implants.

### Tissue section Immunofluorescence, quantification of hydrogel degradation, and immune infiltration

Slides containing tissue sections (10-25 μm thickness) were blocked with 3% normal goat serum (NGS) in 1X PBS + 0.05% Tween-20 (PBS-T). For intracellular antigens 0.2% triton was added to the blocking buffer. Primary antibody dilutions were prepared as follows in 5% NGS in 1X PBST: rat anti mouse CD11b clone M1-70 (BD Pharmingen; #553308) – 1:100, F4/80 clone A3-1 (BioRAD; MCA497G) – 1:400, and IL-33 (abcam; ab187060) – 1:200. Sections were stained with primary antibodies overnight at 4°C, and subsequently washed with 3% NGS in 1X PBS-T. Secondary antibodies (Goat anti-rat Alexa-647, Invitrogen) were all prepared in 5% Normal Goat Serum (NGS) in 1X PBST at a dilution of 1:500. Three 5-minute washes with PBST were performed after each antibody incubation. Sections were incubated in secondary antibodies for 1 hour at room temperature, and subsequently washed with 1X PBST. For multicolor immunofluorescence staining for primary and secondary of each antigen were performed in sequence. Sections were either mounted with antifade mounting medium with DAPI (Fisher Scientific; H1200) or counterstained with 2 μg/ml DAPI in 1X PBST for 30 mins at room temperature and then mounted in Antifade Gold mounting medium.

### Computational analysis of multi-color immunofluorescence images

A MATLAB code was used for analysis of the multicolor immunofluorescence images. The code divides the hydrogel into an Edge region (300um from hydrogel-tissue interface) and a Core region (the center of hydrogel to 200um from the inner boundary of the Edge region). For each hydrogel subregion the code reads CD11b and F4/80 signal, binarizes each to form a mask using a similar threshold for all samples. The code then uses the nuclear stain and IL-33^+^ stains to identify all nuclei and IL-33^+^ cells. The density of each cell type is then quantified by counting the number of nuclei and IL-33^+^ cells overlapping or evading the masks divided by the area of the region of interest. Areas with defects caused by tissue sectioning were excluded from analysis. Although not affecting the code performance, the image condition was kept the same across all samples.

### ELISA

For assessment of anti-L or anti-D antibodies, sera were collected by cardiac puncture 21 days following hydrogel application of mice (subcutaneous implant or application to wound). for detection of anti-L and anti-D antibodies plates were coated with either L-MMP peptide or D-MMP peptide resepcitvely (GenScript; sequence above) Serum samples were tested at a 1:500 dilution followed by incubation with alkaline phosphatase-labeled goat anti-mouse IgG1 or IgG2a, or IgG3 antibodies (Southern Biotechnology Associates or BD Pharmingen), and development with *p*-nitrophenylphosphate substrate (Sigma-Aldrich). Optical density at 405 nm (OD_405_) was read using a spectramax i3X microplate reader (Softmax Pro 3.1 software; Molecular Devices).

### Statistical Analysis

All statistical analysis was performed using Prism 6 (GraphPad, Inc.) software. Specifically, two-tailed t-test or one-way ANOVA were used to determine statistical significance, assuming equal sample variance for each experimental group when comparing individual groups. For ANOVA, post hoc analysis with Tukey multiple comparison. For histological counting, B6 and B6.Rag1^−/−^ sham vs 1:1 L/D-MAP analysis, Wilcoxon signed rank analysis was performed, and B6 vs B6.Rag1^−/−^ and subcutaneous immunofluorescence analysis was performed with t test with Mann-Whitney U test.

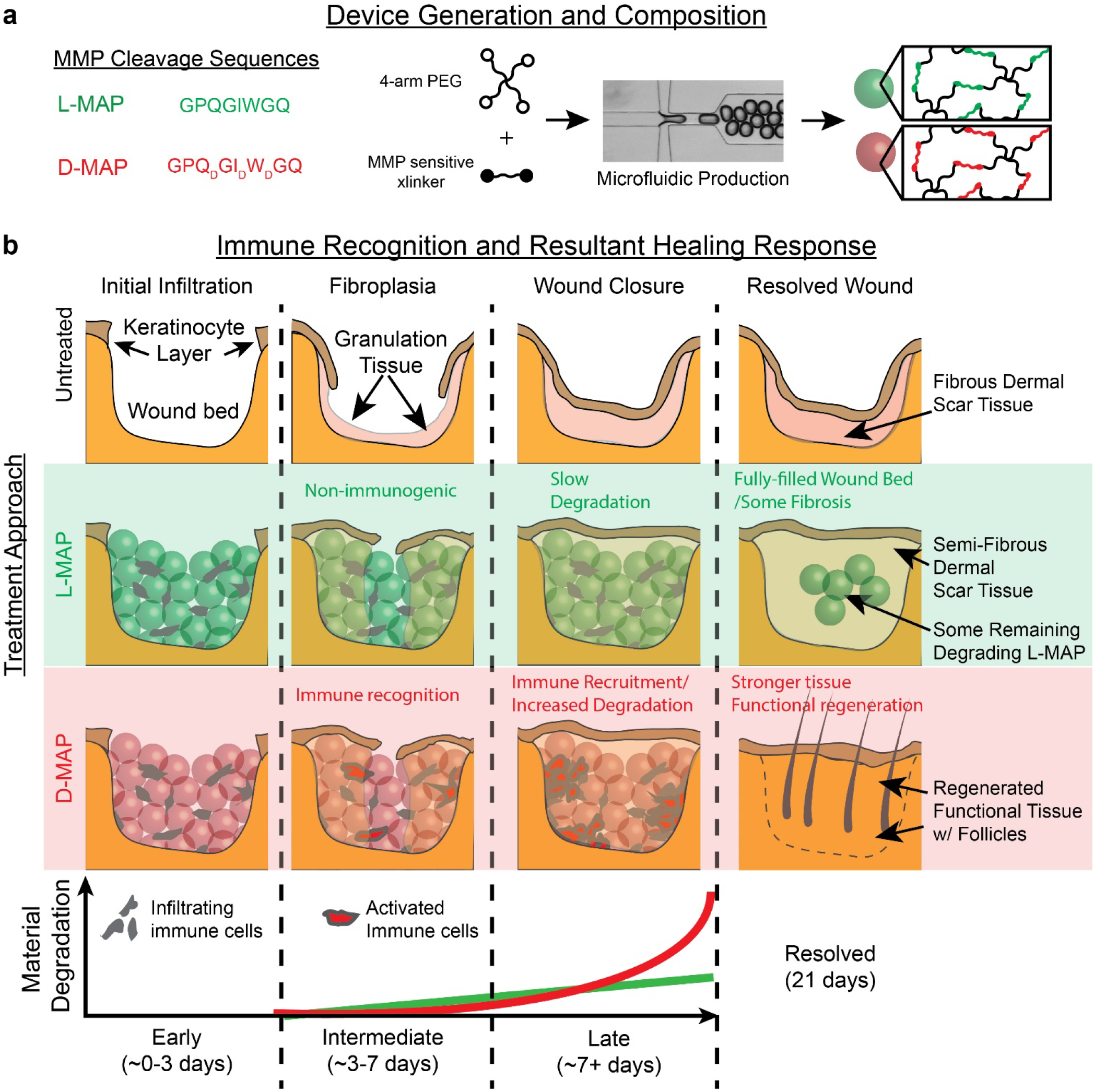
Scheme Figure/Graphical Abstract. **a)** Representation of amino acid chirality within the cross-linking peptides, microfluidic formation of the hydrogel microbeads incorporating L- or D-chirality peptides. **b)** The use of L- or D-MAP in a wound healing model demonstrates that both L- or D-MAP hydrogel fill the wound defect. While wounds that heal in the absence of hydrogel heal with an atrophic scar with loss of tissue, the epidermis forms over the scaffold in both cases and allows for increased dermal thickness. However, in the case of D-MAP, the hydrogel activates the adaptive immune system over time, resulting in tissue remodeling and skin regeneration as the adaptive immune system degrades the D-MAP scaffold.

## Acknowledgements

We would like to thank the National Institutes of Health for funding F32EB018713-01A1 (DRG), T32-GM008042 (MMA), U01AR073159 (MVP), Pew Charitable Trust (MVP), LEO Foundation (MVP), NSF grant DMS1763272 and Simons Foundation Grant (594598, QN) (MVP), R01NS094599 (TS), R01HL110592 (TS), R03AR073940 (POS) and K08AR066545 (POS). We would like to thank Dr. Sasha Cai Lesher-Perez and Michael Bogumil for their assistance with MATLAB coding. We like to thank Yining Liu for assistance running endotoxin texts. We would also like to thank the Advanced Light Microscopy and Spectroscopy at California NanoSystems Institute and Electron Microscopy Core Laboratory of the Brain Research Institute at UCLA and, particularly, for the significant help of Marianne Cilluffo.

## Data Availability

No data sets such as protein, DNA, or RNA sequences, crystallographic data, genetic polymorphisms, linked genotype and phenotype data, macromolecular structure or earth, space and environmental sciences were generated or analyzed during the current study.

## Author Contribution

D.R.G., P.O.S., and T.S. conceived the experiments. D.R.G., W.M.W, E.S., M.M.A., and J.K. carried out microfluidic design and fabrication, and D.D.C. oversaw microfluidic design and fabrication. D.R.G., M.M.A., C.H.K., W.M.W, J.S.W., A.C.F., E.S., A.R., V.R., P.O.S. performed experiments. D.R.G., M.M.A., J.S.W., A.R., M.V.P., T.S. and P.O.S. analyzed and interpreted data. D.R.G., M.M.A., P.O.S., and T.S. wrote the manuscript and all authors discussed the results and contributed to writing portions of the manuscript and editing the manuscript. D.R.G. and M.M.A. contributed equally to this work.

The co-principal investigators are P.O.S. and T.S.

## Supplemental Figure Legend

**Supplemental Figure 1.**
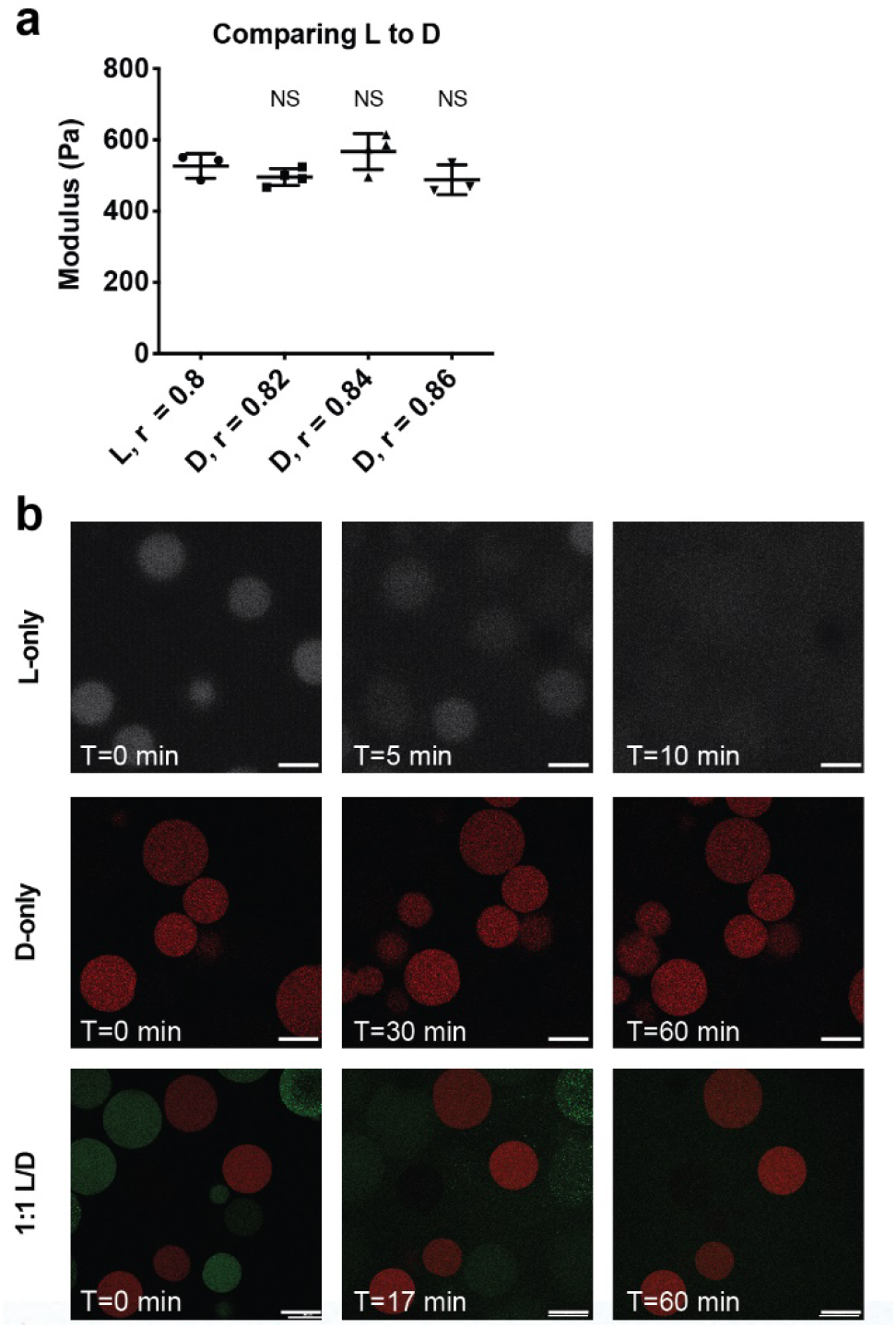
*In vitro* characterization of L- and D-chiral microparticles and MAP hydrogel. **a)** Rheological characterization of MAP hydrogels composed of L or D-peptide crosslinked microgels. The r-ratio (ratio of -SH to -VS) used to form the microgels was changed to arrive at the same storage modulus for both L and D MAP scaffolds. NS represents a no statistical significance between the L MAP scaffold to the D-MAP scaffold indicated using a student t-test. **b)** Collagenase I degradation study of L, D and a 1:1 mixture of L and D microgels. As expected, L-peptide crosslinked microgels are degradable by collagenase I and are completely degraded by 60 minutes. I contrast D-peptide crosslinked microgels have no visible degradation within the 60-minute incubation in collagenase I. In a 1:1 mixture of L and D-peptide crosslinked microgels only L-peptide crosslinked microgels degrade. Images show representative examples of microscope images from *in vitro* hydrogel degradation of L, D, and 1:1 L/D-MAP. Scale = 200μm.

**Supplemental Figure 2.**
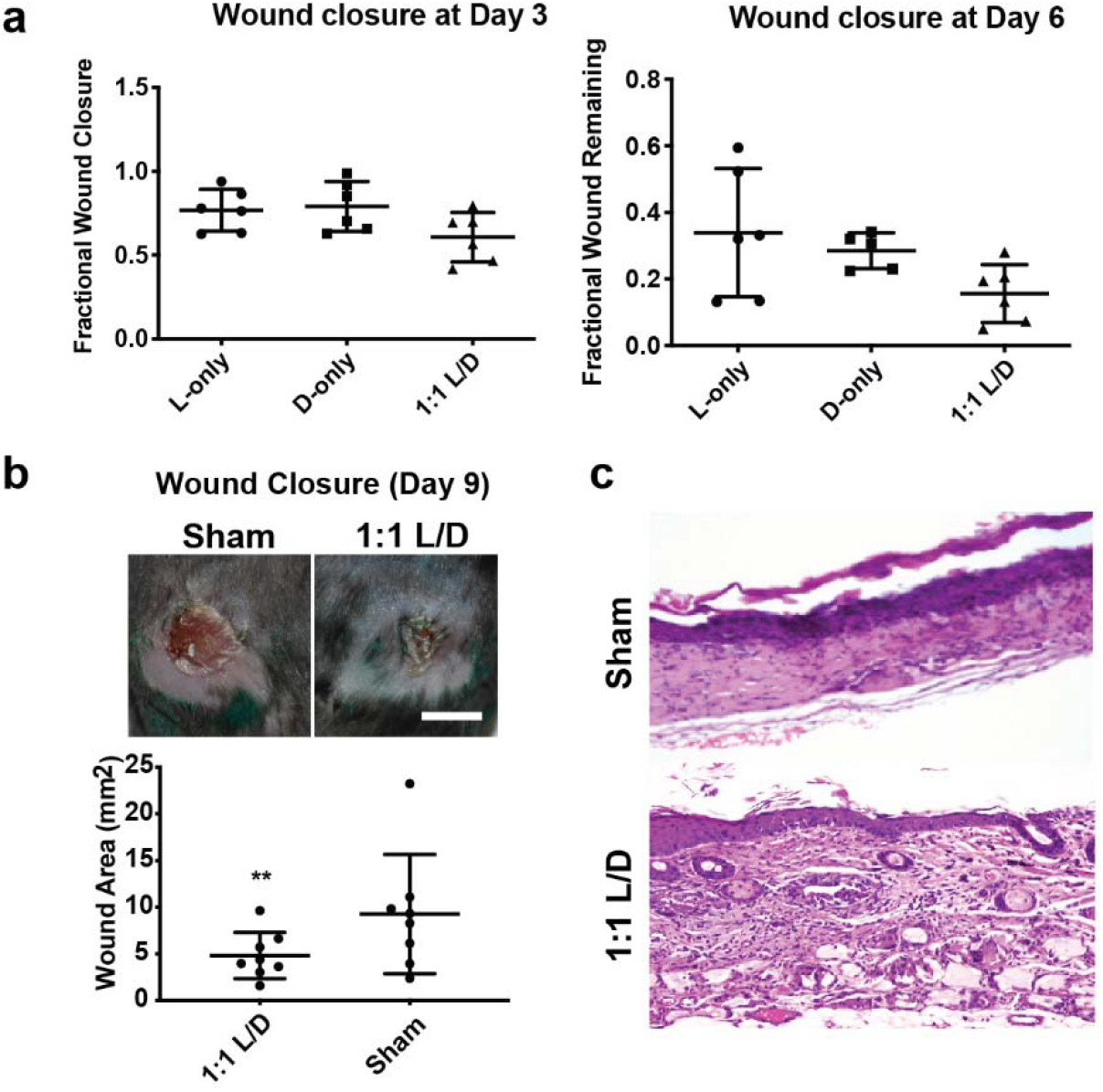
Early wound closure results with different hydrogel treatments. **a)** No difference in wound closure between different L, D, and L/D MAP hydrogel treatment. **b)** Comparison of L and D hydrogel to sham in B6 mice at day 9 reveals improved wound closure when compared to Sham. Scale = 5mm. * denotes p=0.027 by Wilcoxon matched pair signed rank test. C. 100x view of histology from SKH1 (hairless) mice 21 days after wounding demonstrating typical scar formation in sham mice (top) or vellus hair follicles and sebaceous glands directly over several degrading MAP gel particles in a mouse that was treated with D-MAP hydrogel.

**Supplemental Figure 3.**
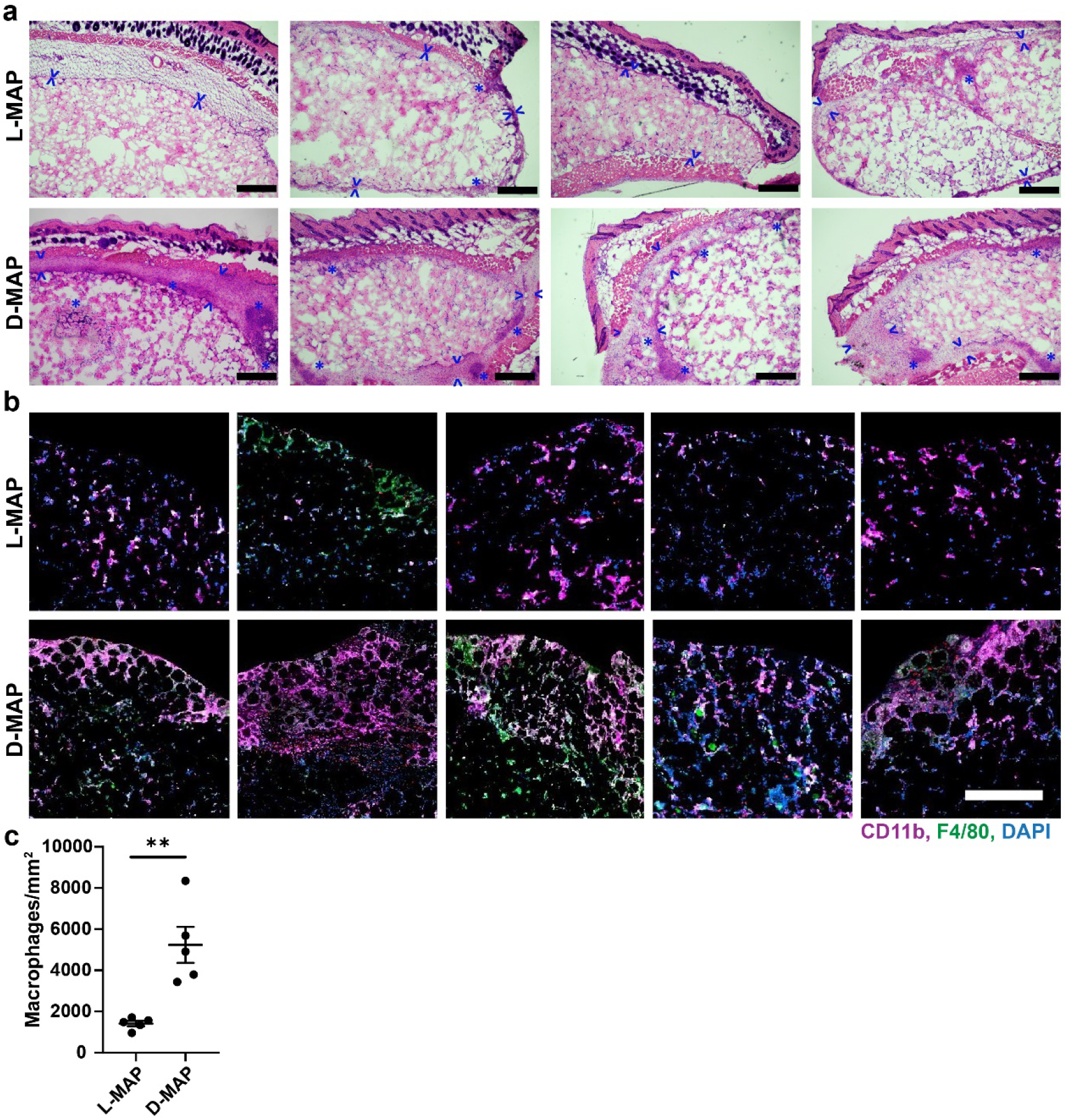
**a)** H&E staining of subcutaneous implants of L-MAP and D-MAP hydrogels. Arrowheads denote the expansion of the fibrous capsule with lymphohistiocytic cells and with admixed neutrophils and eosinophils. Asterisks denote foci of more robust inflammation. Note the minimal fibrous capsule and inflammatory response in L-MAP hydrogels compared to more robust response in D-MAP. Scale = 500μm **b)** Immunofluorescence images from all subcutaneous implants of L-MAP and D-MAP hydrogels for F4/80 (green), CD11b (purple), IL-33 (red), and DAPI (blue). White to light pink denotes co-staining for F4/80 and CD11b antigens. Scale = 500 μm. (a) and (b,c) represent separate experiments. **c)** Quantification of F4/80^+^CD11b^+^ macrophages in the edge of L-MAP and D-MAP implants. ** denotes p=0.0025.

**Supplemental Figure 4.**
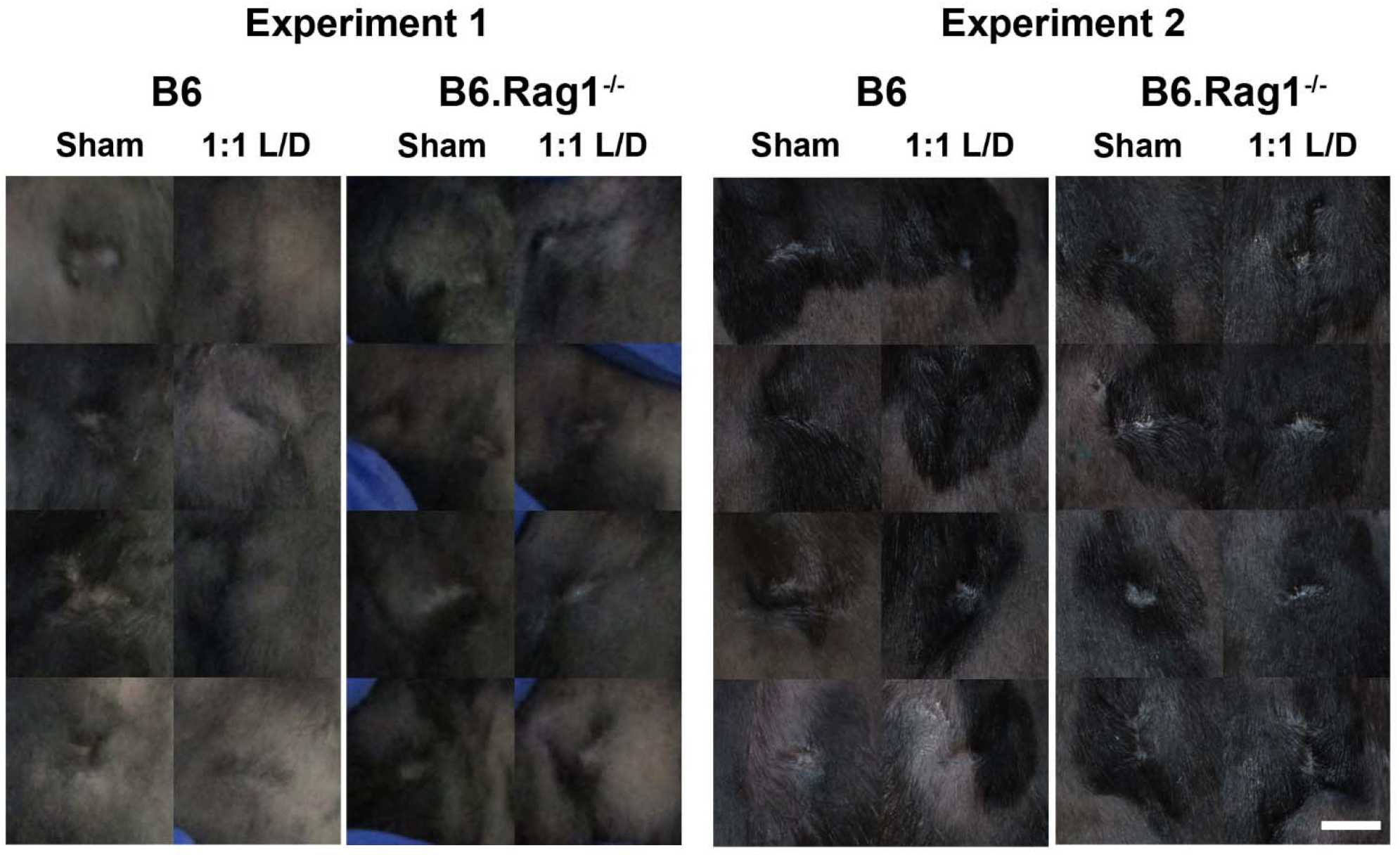
L/D MAP hydrogel diminishes the clinical appearance of scar in WT mice but not B6.Rag1^−/−^ mice. Splinted wounds (6mm) were performed on B6 or B6.Rag1^−/−^ mice, and one side was treated with L/D MAP hydrogel and the other with no hydrogel. On Day 16, clinical photographs of wounds were taken. Shown are all healed wounds, paired by mouse, in two separate experiments. Scale = 2mm.

**Supplemental Figure 5.**
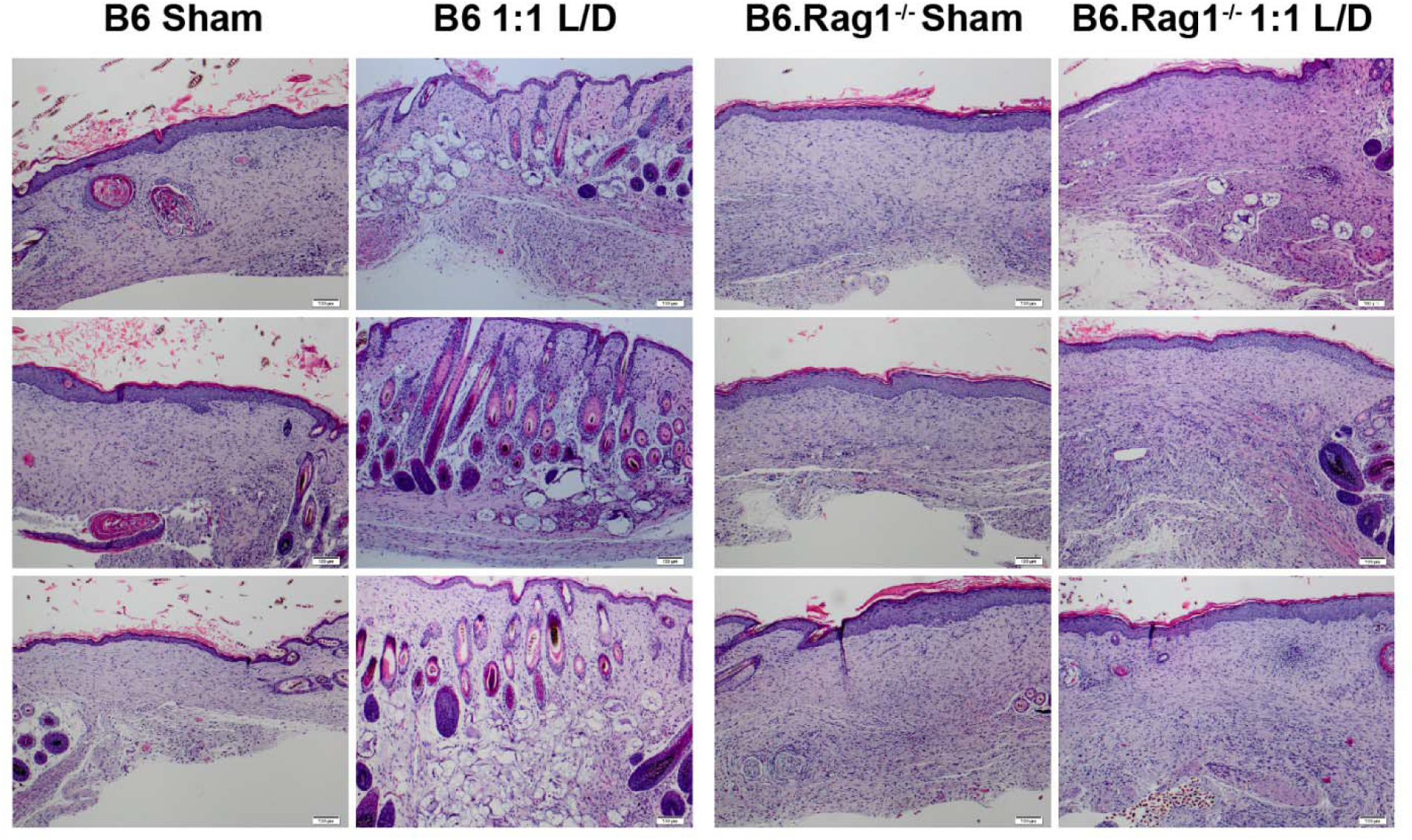
Additional histology from healed 1:1 L/D-MAP or Sham treated wounds in SKH1 (hairless) mice, B6 mice, and B6.Rag1^−/−^ mice. Scale = 100μm. Note in the 1:1 L/D-MAP treated B6 samples, multiple hair follicles and some sebaceous glands, including some in disarrayed orientation compared to the surrounding tissue, are present directly overlying degrading microgels. Similar hair follicles are not present in any other group of samples.

