## Supplemental data for "Activating an adaptive immune response from a hydrogel scaffold imparts regenerative wound healing"

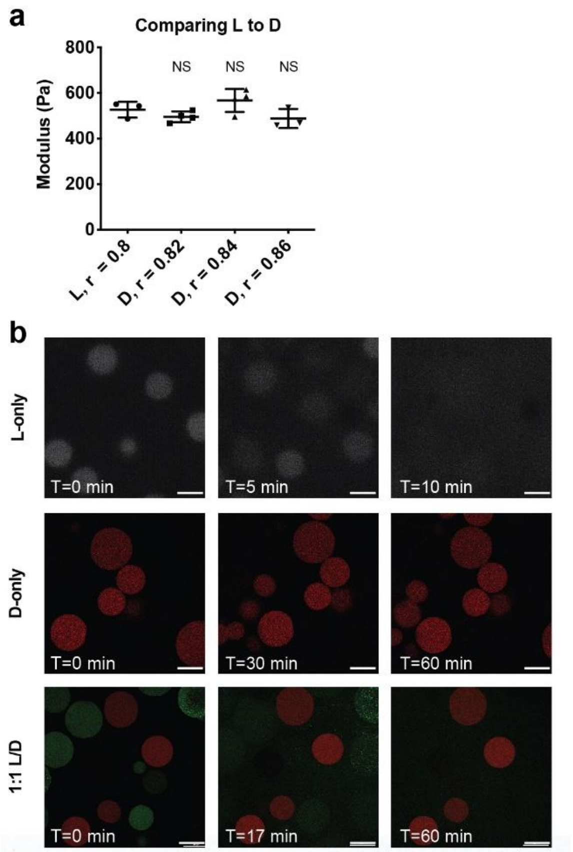


**Supplemental Figure 1.** *In vitro* characterization of L- and D- chiral microparticles and MAP hydrogel. **a)** Rheological characterization of MAP hydrogels composed of L or D-peptide crosslinked microgels. The r-ratio (ratio of -SH to -VS) used to form the microgels was changed to arrive at the same storage modulus for both L and D MAP scaffolds. NS represents a no statistical significance between the L MAP scaffold to the D-MAP scaffold indicated using a student t-test. **b)** Collagenase I degradation study of L, D and a 1:1 mixture of L and D microgels. As expected, L-peptide crosslinked microgels are degradable by collagenase I and are completely degraded by 60 minutes. I contrast D-peptide crosslinked microgels have no visible degradation within the 60-minute incubation in collagenase I. In a 1:1 mixture of L and D-peptide crosslinked microgels only L-peptide crosslinked microgels degrade. Images show representative examples of microscope images from *in vitro* hydrogel degradation of L, D, and 1:1 L/D-MAP. Scale = 200µm.

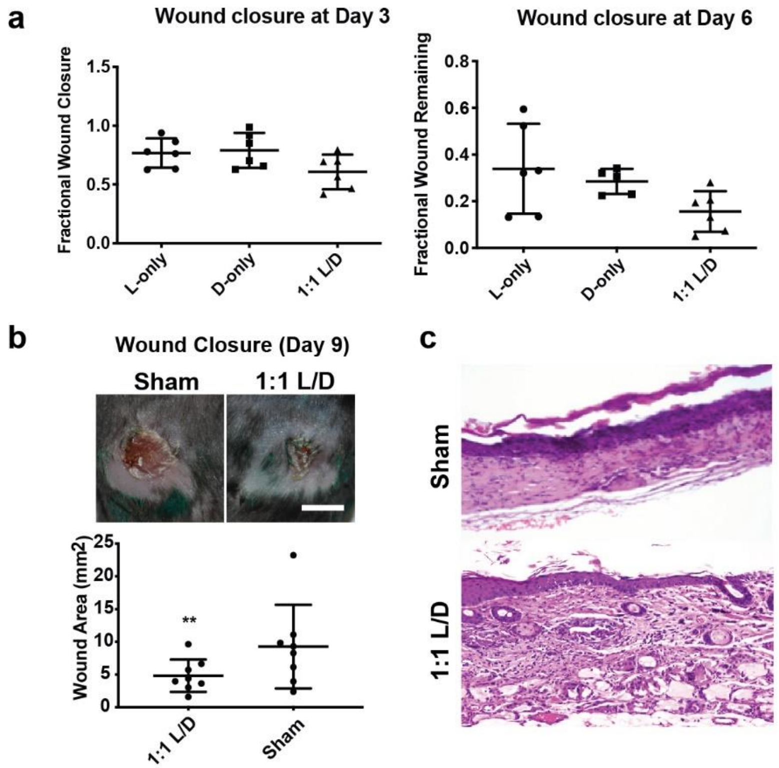


**Supplemental Figure 2.** Early wound closure results with different hydrogel treatments. **a)** No difference in wound closure between different L, D, and L/D MAP hydrogel treatment. **b)** Comparison of L and D hydrogel to sham in B6 mice at day 9 reveals improved wound closure when compared to Sham. Scale = 5mm. * denotes p=0.027 by Wilcoxon matched pair signed rank test. C. 100x view of histology from SKH1 (hairless) mice 21 days after wounding demonstrating typical scar formation in sham mice (top) or vellus hair follicles and sebaceous glands directly over several degrading MAP gel particles in a mouse that was treated with D-MAP hydrogel.

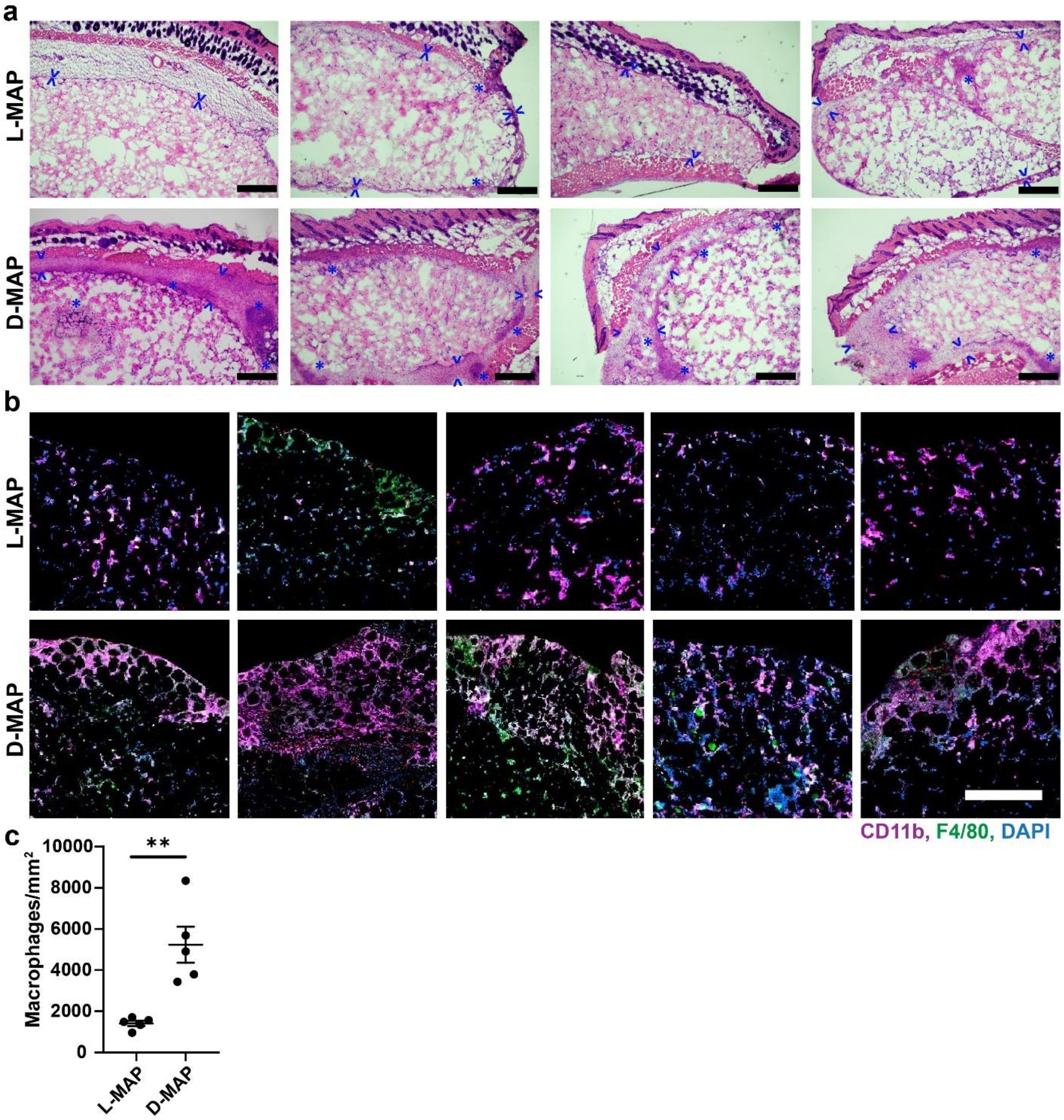


**Supplemental Figure 3. a)** H&E staining of subcutaneous implants of L-MAP and D-MAP hydrogels. Arrowheads denote the expansion of the fibrous capsule with lymphohistiocytic cells and with admixed neutrophils and eosinophils. Asterisks denote foci of more robust inflammation. Note the minimal fibrous capsule and inflammatory response in L-MAP hydrogels compared to more robust response in D-MAP. Scale = 500µm **b)** Immunofluorescence images from all subcutaneous implants of L-MAP and D-MAP hydrogels for F4/80 (green), CD11b (purple), IL-33 (red), and DAPI (blue). White to light pink denotes co-staining for F4/80 and CD11b antigens. Scale = 500 µm. (a) and (b,c) represent separate experiments. **c)** Quantification of F4/80$^+$CD11b$^+$ macrophages in the edge of L-MAP and D-MAP implants. ** denotes p=0.0025.

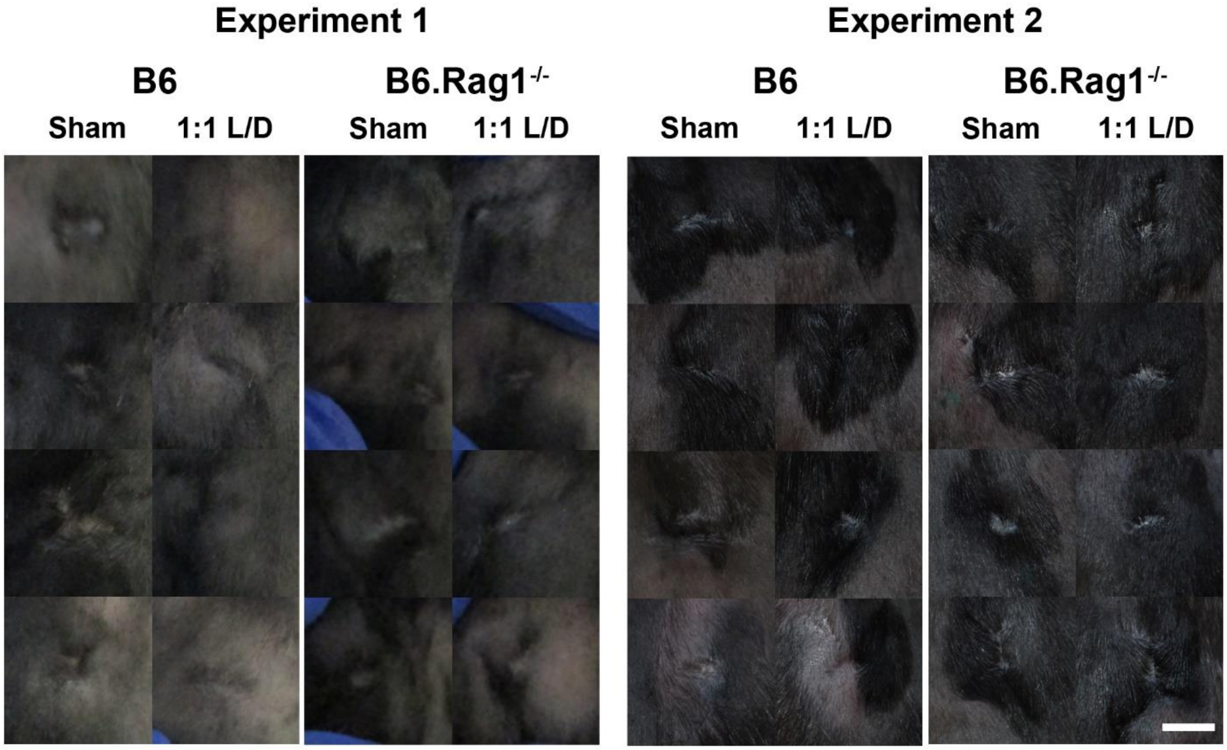


**Supplemental Figure 4.** L/D MAP hydrogel diminishes the clinical appearance of scar in WT mice but not B6.Rag1$^{-/-}$ mice. Splinted wounds (6mm) were performed on B6 or B6.Rag1$^{-/-}$ mice, and one side was treated with L/D MAP hydrogel and the other with no hydrogel. On Day 16, clinical photographs of wounds were taken. Shown are all healed wounds, paired by mouse, in two separate experiments. Scale = 2mm.

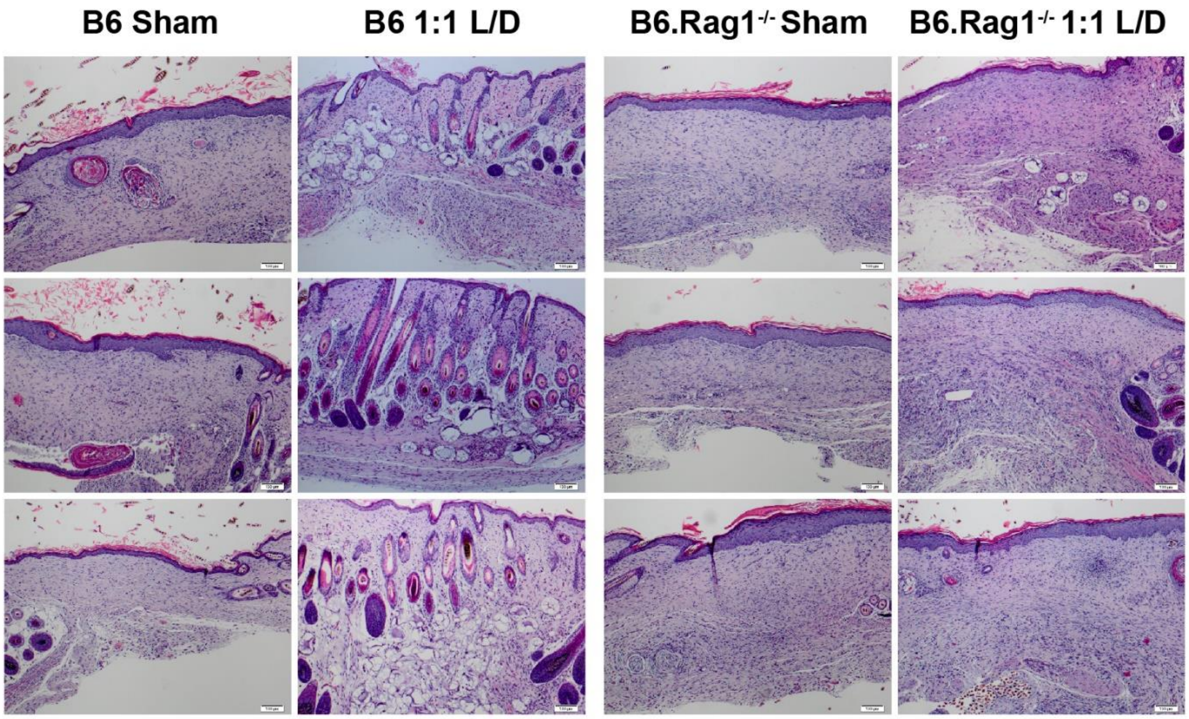


**Supplemental Figure 5.** Additional histology from healed 1:1 L/D-MAP or Sham treated wounds in SKH1 (hairless) mice, B6 mice, and B6.Rag1$^{-/-}$ mice. Scale = 100µm. Note in the 1:1 L/D-MAP treated B6 samples, multiple hair follicles and some sebaceous glands, including some in disarrayed orientation compared to the surrounding tissue, are present directly overlying degrading microgels. Similar hair follicles are not present in any other group of samples.
